# Glia generate distinct visual processing centres by locally inhibiting ERK activity in an optic lobe neuroepithelium

**DOI:** 10.64898/2026.04.06.716791

**Authors:** Bradleigh M. J. Cocker, Matthew P. Bostock, Haoruo Wei, Vilaiwan M. Fernandes

## Abstract

Epithelial patterning is fundamental to organ development. Extensive work has focused on how neuroepithelia are patterned to generate diverse progenitors, yet how a single neuroepithelium is partitioned to produce distinct processing centres is poorly understood. Here, we focus on the Drosophila outer proliferation centre neuroepithelium, which generates the medulla and lamina visual processing centres. Medulla neuroblasts are produced by a proneural wave that initiates at the medial margin of the neuroepithelium and moves laterally propagated by EGFR-ERK signalling, whereas lamina precursors arise at the lateral neuroepithelial margin and have been proposed to require photoreceptor-derived Hedgehog. Here, we show that Hedgehog signalling is dispensable for lamina precursor specification but instead promotes their survival. In contrast, suppressing ERK and apoptosis together is sufficient to drive ectopic lamina precursor development. We find that cortex glia secrete the EGF antagonist Argos, which accumulates at the lateral neuroepithelium, thus repressing ERK activity locally. Together, our findings reveal a glia-mediated, extrinsic patterning mechanism that suppresses EGFR-ERK signalling in the lateral neuroepithelium, protecting these cells from the proneural wave and instructing lamina over medulla fate.

**Summary statement:** Glial cells locally control ERK signalling to partition a developing tissue, enabling distinct structures to arise from a common neuroepithelium.

## Introduction

Epithelial patterning is fundamental to organ development, as it generates distinct cell types in appropriate locations to support tissue architecture and function. In developing nervous systems, much work has focused on how neuroepithelia are patterned into discrete spatial domains by morphogens and gene regulatory networks, establishing distinct progenitor identities that in turn generate the diverse neuronal populations of individual processing centres (Cohen et al., 2013; Li et al., 2013; Erclik et al., 2017; Sagner and Briscoe, 2019; Konstantinides et al., 2022). A distinct and relatively underexplored axis of neuroepithelial patterning relates to how a single neuroepithelium is subdivided to generate distinct processing centres. To address this question, we focussed on the genetically tractable *Drosophila melanogaster* visual system, where two distinct visual processing centres arise from a common neuroepithelium.

The Drosophila visual system is composed of the compound eyes and optic lobes. Each optic lobe contains four distinct visual processing centres (neuropils), the lamina, medulla, lobula and lobula plate, which are built during the third larval instar (L3) and pupal stages. The lamina and medulla arise from a crescent-shaped neuroepithelium at the surface of the optic lobe, called the outer proliferation centre (OPC; Fig. 1A,B), whereas the lobula and lobula plate originate from the inner proliferation centre neuroepithelium, located deeper in the optic lobe. Here, we focus on the OPC neuroepithelium and how it is subdivided to produce the lamina and medulla. These structures form retinotopic maps with the compound eye, yet differ markedly in their neurogenic programs and cellular complexity: post-mitotic lamina precursor cells differentiate into just 5 neuron types in response to extrinsic signals (Apitz and Salecker, 2014; Bostock et al., 2022; Prasad et al., 2022; Donoghue et al., 2025), whereas medulla neuroblasts undergo numerous rounds of asymmetric divisions, are subject to spatial and temporal patterning and generate ∼100 neuronal types (Fischbach and Dittrich, 1989; Li et al., 2013; Bertet et al., 2014; Erclik et al., 2017; Ozel et al., 2021). Thus, it is striking that these two structures, with such distinct neurogenic programmes and outputs, arise from the same neuroepithelium.

**Fig. 1:**
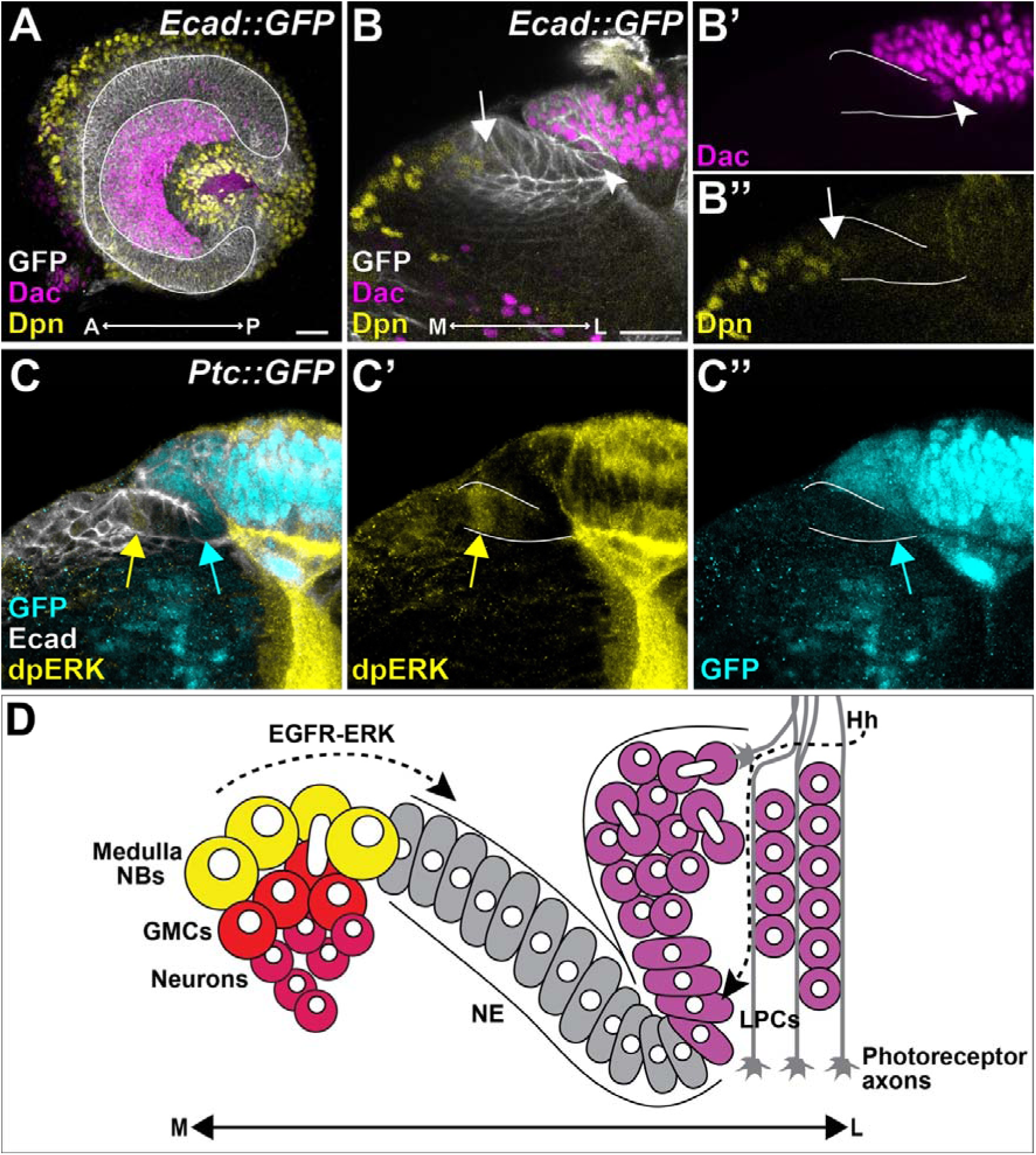
ERK and Hh signalling are regionalised in the OPC neuroepithelium. **(A-B)** Representative optic lobes from late L3 larvae expressing Ecad::GFP (grey), which marks the OPC neuroepithelium. Dac (magenta) marks lamina precursors and lamina neurons and Dpn (yellow) marks neuroblasts (N=10). (A) A maximum projection showing the lateral view of the optic lobe with the crescent shaped neuroepithelium. (B) A cross-sectional view shows the lamina furrow. White arrow marks Dpn expression in the youngest medulla neuroblast, white arrowhead marks the onset of Dac expression in the neuroepithelium. **(C)** Representative optic lobe from a late L3 larva expressing Ptc::GFP (cyan), a target of Hh signalling. Columnar cells of the neuroepithelium are marked by Ecad (grey). dpERK (yellow) cells with high ERK signalling (N=3). Yellow arrow marks dpERK expression in the medial neuroepithelium, cyan arrow marks the onset of Ptc::GFP expression in the neuroepithelium. **(D)** Schematic of the OPC neuroepithelium partitioned into medulla and lamina lineages. A wave of EGFR-ERK signalling, which moves laterally from the medial margin drives medulla neuroblasts formation, whereas Hh from photoreceptor axons at the lateral neuroepithelium is required for lamina precursor differentiation. White lines indicate the neuroepithelium, scale bar = 20 μm. Anterior (A), posterior (P), medial (M), lateral (L), neuroblasts (NBs), ganglion mother cells (GMCs), neuroepithelial cells (NE), lamina precursor cells (LPCs).

Early in L3, a proneural wave initiates at the medial margin of the OPC neuroepithelium, sequentially converting neuroepithelial cells into medulla neuroblasts marked by Deadpan (Dpn) expression (Fig. 1A,B) (Yasugi et al., 2008; Egger et al., 2010; Yasugi et al., 2010; Sato et al., 2016; Jörg et al., 2019). Epidermal Growth Factor Receptor - Extracellular Signal-Regulated Kinase (EGFR-ERK) signalling drives proneural wave progression in the medial neuroepithelium (Fig. 1C,D), counterbalanced by Jak/Stat and Notch signalling, which regulate the pace of progression (Yasugi et al., 2008; Egger et al., 2010; Yasugi et al., 2010; Sato et al., 2016; Jörg et al., 2019). In parallel, at the lateral side, the OPC neuroepithelium folds into a deep groove known as the lamina furrow (Fig. 1C,D), with cells arrested in the G1 phase of the cell cycle within the lateral furrow (Selleck et al., 1992). Photoreceptors born in the developing eye disc extend axons through the optic stalk and into the optic lobe where they contact the neuroepithelium at the trough of the lamina furrow (Selleck et al., 1992; Huang and Kunes, 1996; Huang and Kunes, 1998). These axons secrete Hedgehog (Hh), activating Hh signalling in neuroepithelial cells in the lateral half of the furrow (Figure 1C,D) (Huang and Kunes, 1996; Huang and Kunes, 1998). Here, Hh signalling has been proposed to drive lamina development by inducing expression of the lamina precursor cell (LPC) marker Dachshund (Dac) in neuroepithelial cells of the lateral furrow, followed by delamination from the neuroepithelium, cell cycle progression and terminal divisions, and assembly into columns along photoreceptor axons (Huang and Kunes, 1996; Huang and Kunes, 1998).

At first glance, these spatially restricted signals appear sufficient to partition the neuroepithelium into medial and lateral domains, with EGFR-ERK activity promoting medulla neuroblast specification medially and photoreceptor-derived Hh promoting lamina precursor formation at the lateral furrow (Fig. 1C,D). Consistent with this view, hyperactivating EGFR-ERK signalling in the neuroepithelium drives ectopic medulla neuroblast formation and disrupts lamina development (Yasugi et al., 2010). However, confoundingly, lamina development is also disrupted by loss of EGFR signalling (Yasugi et al., 2010), and in contrast, experiments hyperactivating Hh signalling in the absence of photoreceptor axons rescue lamina development, but only lateral to the lamina furrow and do not induce ectopic lamina medially (Huang and Kunes, 1996; Huang and Kunes, 1998). This spatially restricted response to Hh signalling has been attributed to cell-cycle-dependent competence gating within the lamina furrow such that only cells arrested in G1 are thought to be competent to respond to Hh (Huang and Kunes, 1998) and therefore imply that other processes must be at play even if only to restrict cell-cycle progression and competence to respond to Hh.

To determine how the neuroepithelium is partitioned into lamina and medulla, we re-examined the role of Hh signalling in lamina development. We found that Hh signalling acts permissively to promote lamina precursor survival but that it is dispensable for initiating Dac expression and for cell cycle progression through lamina precursor terminal divisions. In contrast, suppressing ERK signalling, in combination with blocking apoptosis, was sufficient to induce ectopic lamina precursors from the medial neuroepithelium. Consistent with this, we observed that ERK activity is locally suppressed in the lamina furrow and that this suppression is required for lamina precursor specification. We further show that cortex glia overlying the neuroepithelium secrete Argos (Aos), an antagonist of the EGF Spitz, which accumulates at high levels within the trough of the lamina furrow. The lamina fails to develop in *aos* mutants, but restoring *aos* expression specifically in cortex glia rescues lamina formation. Together, these findings reveal a glia-mediated, extrinsic patterning mechanism that locally suppresses EGFR signalling in the lamina furrow, protecting these cells from being consumed by the proneural wave and instructing lamina over medulla fate.

## Results

### Hh signalling is not sufficient to induce ectopic lamina fate throughout the neuroepithelium

The sufficiency of Hh signalling to induce lamina development has primarily been evaluated in genetic backgrounds lacking photoreceptor axons (Huang and Kunes, 1996; Huang and Kunes, 1998). In these contexts, Hh signalling was shown to be sufficient to induce lamina formation lateral to the lamina furrow but not medial to it (Huang and Kunes, 1996; Huang and Kunes, 1998), leaving unresolved the extent to which Hh signalling can induce lamina fate more broadly. To address this, we tested whether cell-autonomous hyperactivation of the Hh pathway was sufficient to induce ectopic lamina in a wild-type context. *patched* (*ptc*) encodes a transcriptional target and negative regulator of Hh signalling (Chen and Struhl, 1996; Chen and Struhl, 1998). We generated GFP-labelled clones mutant for *ptc* (*ptc^S2^;* Phillips et al., 1990) using the Mosaic Analysis with a Repressible Cell Marker (MARCM) technique (Lee and Luo, 2001).

In wild-type optic lobes, Dac expression initiated within the neuroepithelium in the lateral half of the lamina furrow (Fig. 2A). In *ptc^S2^* clones spanning the lamina furrow, Dac expression displayed a mild medial expansion, initiating ∼3 cells medial to the lamina trough (Fig. 2B,C). However, despite premature Dac expression within the trough of the lamina furrow, cells at the medial margins of the neuroepithelium never expressed Dac even when encompassed by *ptc^S2^* clones (0/25 clones). Moreover, *ptc^S2^*clones contributed to medulla lineages as streams of cells, indicating that *ptc^S2^* clones arising in the medial neuroepithelium differentiated into medulla neuroblasts, capable of generating ganglion mother cells (GMCs) and neurons (Fig. 2B, arrowhead). Together, these results show that Hh signalling is not sufficient to induce lamina fate throughout the neuroepithelium. This could reflect a restriction in competence, whereby medial neuroepithelial cells fail to respond to Hh, as suggested previously (Huang and Kunes, 1998), or alternatively that Hh signalling acts permissively rather than instructively during lamina development and other signals are required to instruct lamina fate. To distinguish between these possibilities, we next re-examined the requirement for Hh signalling in specifying lamina fate.

**Fig. 2:**
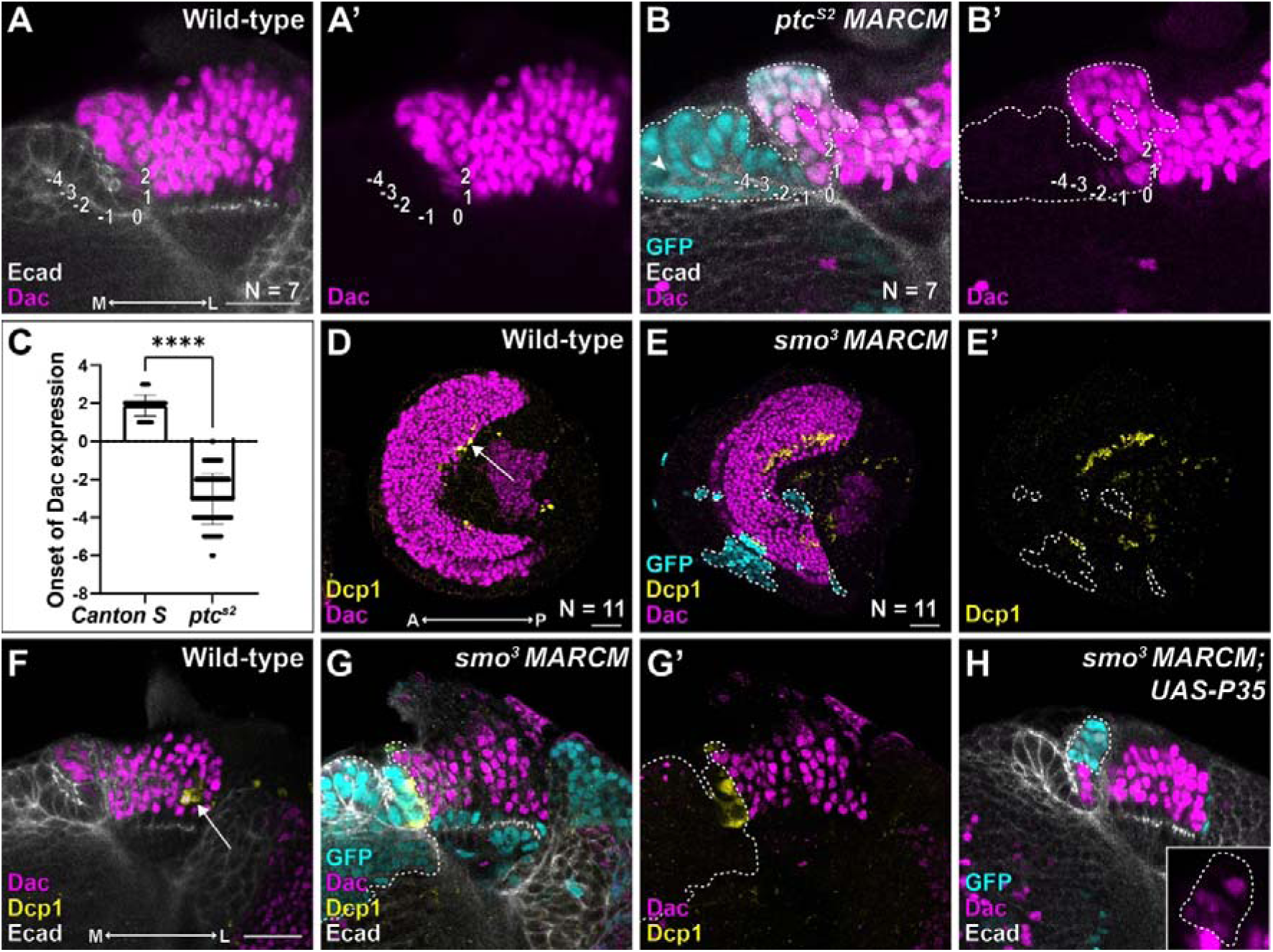
Hh signalling is insufficient to induce ectopic lamina precursors in the medial neuroepithelium and is necessary for lamina precursor survival but dispensable for their specification. **(A,B)** Representative cross-sections of wild-type optic lobes (A; N=7 lobes) and optic lobes containing *ptc^S2^* MARCM clones (B; N=7 lobes) marked by GFP (cyan - outlined) stained for Ecad (grey) and Dac (magenta). Numbers indicate cell positions within the lamina furrow, where 0 is the trough of the lamina furrow. **(C)** Quantification of the onset of Dac expression in the wild-type and *ptc^S2^* MARCM clones (Mann Whitney U test, P<0.0001). **(D,E)** Representative lateral views of a wild-type optic lobe (D; N=11) and an optic lobe containing *smo^3^* MARCM clones marked by GFP (cyan - outline) stained for Dac (magenta) and Dcp1 (yellow). Arrow (D) indicates normal regions of apoptosis in the lamina. **(F-H)** Representative cross-sections of a wild-type optic lobe (F) and optic lobes with *smo^3^*MARCM clones (G; cyan – outline) and *smo^3^* MARCM clones expressing P35 (H; cyan – outline) stained for Dac (magenta), Ecad (grey), and Dcp1 (F,G yellow). All optic lobes are from pupae dissected at 0h APF. Scale bar = 20 μm. Anterior (A), posterior (P), medial (M), lateral (L).

### Hh signalling is required for lamina precursor survival but dispensable for their specification and cell cycle progression

We disrupted Hh signalling cell autonomously by generating MARCM clones homozygous for a mutant allele of the Hh signalling transducer *smoothened*, (*smo^3^*) (Chen and Struhl, 1996). Control clones were recovered throughout the lamina, where they contained Dac expressing cells (Fig. S1A). In contrast, *smo^3^* clones spanning the lamina furrow disrupted lamina development, with disrupted Dac expression in all *smo^3^* clones (Figs. S1B). Interestingly, 65% of *smo^3^* clones lacked Dac expression completely (Fig. S1B; asterisks), whereas 35% of clones retained a small number of cells expressing reduced levels of Dac (Fig. S1B; arrowhead). Notably, *smo^3^* clones retaining Dac expression were restricted to the tips of the lamina crescent (Fig. S1B,C). Crucially, clone induction positions did not differ between control and *smo^3^* clones (Fig. S1D), indicating that the variability in Dac expression does not arise from differences in clone induction between the control and *smo^3^*mutant.

Together, these data indicate that while most of the lamina requires Hh signalling for Dac expression, this requirement is not uniform across the tissue, as Dac expression can be retained at the tips of the lamina crescent. This spatial heterogeneity suggests that Hh signalling may not act strictly instructively, but instead could play a more permissive role in lamina development.

To explore this possibility, we examined *smo^3^* clones in greater detail to determine why lamina precursors fail to develop in the absence of Hh signalling. Previous work has shown that Hh signalling promotes survival cell autonomously in the Drosophila wing pouch (Lu et al., 2017). Therefore, we assessed apoptosis by examining cleaved Death caspase-1 (Dcp1) expression in *smo^3^* clones. In wild-type optic lobes, a small fraction of cells that have incorporated into lamina columns undergoes apoptosis (Prasad et al., 2022), however *smo^3^* clones spanning the lamina furrow exhibited ectopic Dcp1 expression within the lateral lamina furrow compared to wild-type optic lobes, indicating increased apoptosis during lamina precursor specification in the absence of Hh signalling (Fig. 2D-G).

To determine whether Hh signalling instructs lamina precursor development or instead acts permissively to promote survival of already specified lamina precursors, we blocked apoptosis in *smo^3^* clones by expressing the baculovirus caspase inhibitor P35. Strikingly, we found that Dac expression was partially rescued in 74% of *smo^3^* clones expressing P35 (Figs. 2H and S1E), a significant increase compared to 35% *smo^3^* clones (Figs. 2E,G and S1B,E; Fisher’s Exact test, p<0.0002). Furthermore, Dac+ *smo^3^* clones expressing P35 were recovered across the entire lamina, whereas Dac^+^ *smo^3^* clones were restricted to the lamina tips (Fig. S1C,F). Notably, Dac expression levels in *smo^3^* clones expressing P35 were reduced compared to wild-type, suggesting that while Hh signalling is dispensable for initiating Dac expression, it may enhance or stabilise Dac expression in lamina precursors.

In addition to its proposed role in inducing Dac expression, Hh signalling has been implicated in promoting cell cycle progression and terminal divisions (Huang and Kunes, 1996; Huang and Kunes, 1998). To determine whether Hh also acts permissively in this context by promoting cell survival, we examined whether *smo^3^* clones expressing P35 could progress through S-phase and mitosis. Using a brief pulse of 5-ethynyl-2′-deoxyuridine (EdU) to label cells in S-phase and phospho-histone H3 (pH3) to detect mitotic cells, we found that Dac+ lamina precursors in *smo^3^* clones expressing P35 incorporated EdU and were pH3-positive, comparable to the wild-type (Fig. S1G-J). These data indicate that, when rescued from apoptosis, lamina precursors lacking Hh signalling can progress through the cell cycle.

Altogether, these results show that Hh signalling acts permissively to promote lamina precursor survival but is dispensable for instructing their specification or cell cycle progression.

### ERK signalling promotes medulla neuroblast formation while antagonising lamina development

Having established that Hh signalling acts permissively rather than instructively in lamina development, we next sought to identify the signals that instruct lamina fate, and in particular, address how neuroepithelial cells are spatially segregated along the medial-lateral axis to adopt medulla or lamina fates, respectively. EGFR-ERK signalling is required for proneural wave progression and medulla neuroblast formation, and elevated ERK activity has been reported to disrupt lamina development (Yasugi et al., 2008; Egger et al., 2010; Yasugi et al., 2010; Sato et al., 2016; Jörg et al., 2019), suggesting that it may promote medulla fate at the expense of lamina fate. However, previous studies have also reported disrupted lamina development under conditions of both reduced EGFR and increased EGFR signalling, such as in the case of EGFR hypomorphic mutants for loss-of-function, or mutants for *argos (aos),* which encodes a secreted antagonist of the EGF Spitz and overexpression of the transcriptional effector Pointed-P2 (PntP2) for gain-of-function (Yasugi et al., 2010; Morante et al., 2013). These effects have been attributed to indirect roles for EGFR-ERK in maintaining or expanding the neuroepithelium through proliferation, rather than to direct effects on lamina specification (Yasugi et al., 2010; Morante et al., 2013).

To test whether EGFR-ERK signalling antagonises lamina development in addition to promoting medulla neuroblast formation, we increased EGFR-ERK activity within a spatially restricted region of the neuroepithelium. We used pxb^ts^-Gal4 (pxb-Gal4 combined with a temperature sensitive Gal80) to mis-express a constitutively active form of EGFR (EGFR^ACT^) in the central neuroepithelium for 48 hours from early L3 (Queenan et al., 1997; Suzuki et al., 2016a; Courgeon and Desplan, 2019). In contrast to control optic lobes expressing CD8::GFP, expression of EGFR^ACT^ led to the absence of Dac-expressing lamina precursors and neurons in the pxb^ts^-Gal4 domain; instead, the neuroblast marker Dpn was expressed ectopically throughout the neuroepithelium within the pxb^ts^-Gal4 domain (Fig. 3A-F). Consistent with this, RNAi-mediated knockdown of *anterior open* (*aop*; O’Neill et al., 1994), which encodes a repressor of ERK-dependent transcription, driven by pxb^ts^-Gal4 produced the same phenotype, with loss of Dac expression and ectopic Dpn induction (Fig. 3G-I).

**Fig. 3:**
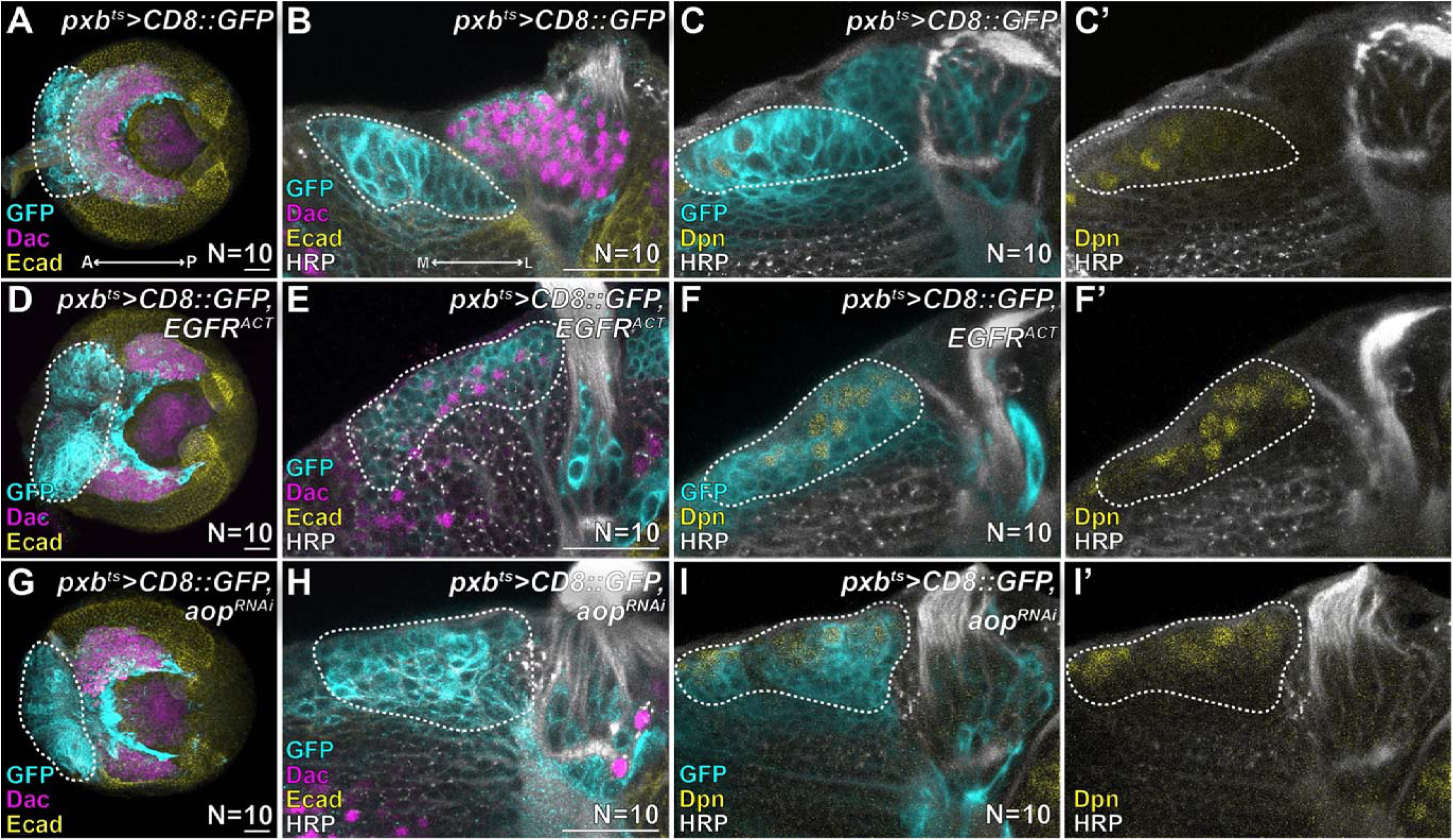
ERK signalling promotes medulla neuroblasts formation at the expense of the lamina. Representative images of Late L3 optic lobes with **(A-C)** pxb-Gal4^ts^ driving expression of CD8::GFP **(D-F)** pxb-Gal4^ts^ driving expression of CD8::GFP and EGFR^ACT^ **(G-I)** pxb-Gal4^ts^ driving expression of CD8::GFP and aop^RNAi^. (A,B, D, E, G and H) show GFP (cyan), Dac (magenta), Ecad (yellow) and horseradish peroxidase (HRP; grey) which stains axons (N=10 lobes each). (C,F and I) show Dpn (yellow) and HRP (grey) (N=10 lobes each). (A,D and G) are maximum projections of optic lobes oriented laterally, (B,C,E,F,H,I) are cross-sectional views. The pxb expression domain of the neuroepithelium is outlined in each image. Scale bar = 20 μm. Anterior (A), posterior (P), medial (M), lateral (L).

Aop is expressed throughout the neuroepithelium, but its localisation appears more nuclear in the trough of the lamina furrow and into the lateral furrow (Fig. S2A) (Weng et al., 2012). To test whether lamina development requires cell-autonomous repression of ERK signalling targets, we generated MARCM clones homozygous for a mutant allele of *aop* (*aop^XE18^*) (Karim et al., 1996; Sawyer et al., 2020). Whereas control clones contributed to both medulla and lamina regions (Fig. S2B), *aop^XE18^* mutant clones were excluded from the lamina and instead recovered within the medulla and other neuropils (Fig. S2C). Together, these results suggest that, in addition to promoting medulla neuroblast formation, EGFR-ERK signalling also antagonises lamina development.

### Inhibiting both ERK signalling and apoptosis in the neuroepithelium drives ectopic lamina precursor development

Given that EGFR-ERK signalling antagonises lamina development, we tested whether suppressing it was sufficient to drive lamina fate. EGFR-ERK signalling is known to be required for cell survival in other contexts in Drosophila (Bergmann et al., 1998; Domínguez et al., 1998; Kurada and White, 1998; Bergmann et al., 2002; Brown and Freeman, 2003; Crossman et al., 2018). Therefore, we blocked apoptosis when disrupting ERK signalling clonally to increase clone recovery.

We first generated MARCM clones mutant for *Death regulator Nedd2-like caspase* (*Dronc^I24^*) to block apoptosis and expressed a constitutively active form of Aop (Aop^ACT^) to repress transcription downstream of ERK signalling in these clones (Rebay and Rubin, 1995). We recovered large control *Dronc^I24^* clones within the medulla cortex. Cells within these clones expressed the pan-neuronal marker Embryonic lethal abnormal visual system (Elav) (Robinow and White, 1991), indicating that they had differentiated into medulla neurons. In contrast, numerous small *Dronc^I24^* clones expressing Aop^ACT^ were recovered within the medulla cortex but lacked Elav expression and instead displayed low levels of Dac expression (Fig. 4B; inset in B’). Although a subset of medulla neurons are Dac+, they also express Elav (Courgeon and Desplan, 2019); therefore, the absence of Elav in *Dronc^I24^*clones expressing Aop^ACT^ indicates that these Dac+ cells are not medulla neurons. Instead, we hypothesised that these clones represent cells that have delaminated from the neuroepithelium and begun differentiating into lamina precursors. E-cadherin (Ecad) and Dac are co-expressed in early lamina precursors (Fig. 1B), therefore we next tested their expression in these clones. Whereas control *Dronc^I24^* clones medial to the lamina furrow did not express ectopic Ecad or Dac, *Dronc^I24^* clones expressing Aop^ACT^ recovered medial to the lamina furrow or in the medulla cortex co-expressed Ecad and Dac, with Ecad often localised in rosette-like structures (Fig. 4C,D), consistent with our hypothesis. As an independent approach, we generated clones homozygous for an amorphic allele of *pnt* (*pnt*^Δ*88*^) (Brunner et al., 1994) and blocked apoptosis by expressing P35. Again, we observed clones medial to the lamina furrow and in the medulla cortex that co-expressed Dac and Ecad specifically in *pnt*^Δ*88*^ clones expressing P35 but not in control clones expressing P35 alone (Fig. 4E,F), suggesting that these cells originated in the medial neuroepithelium, delaminated and initiated differentiation into lamina precursors.

**Fig. 4:**
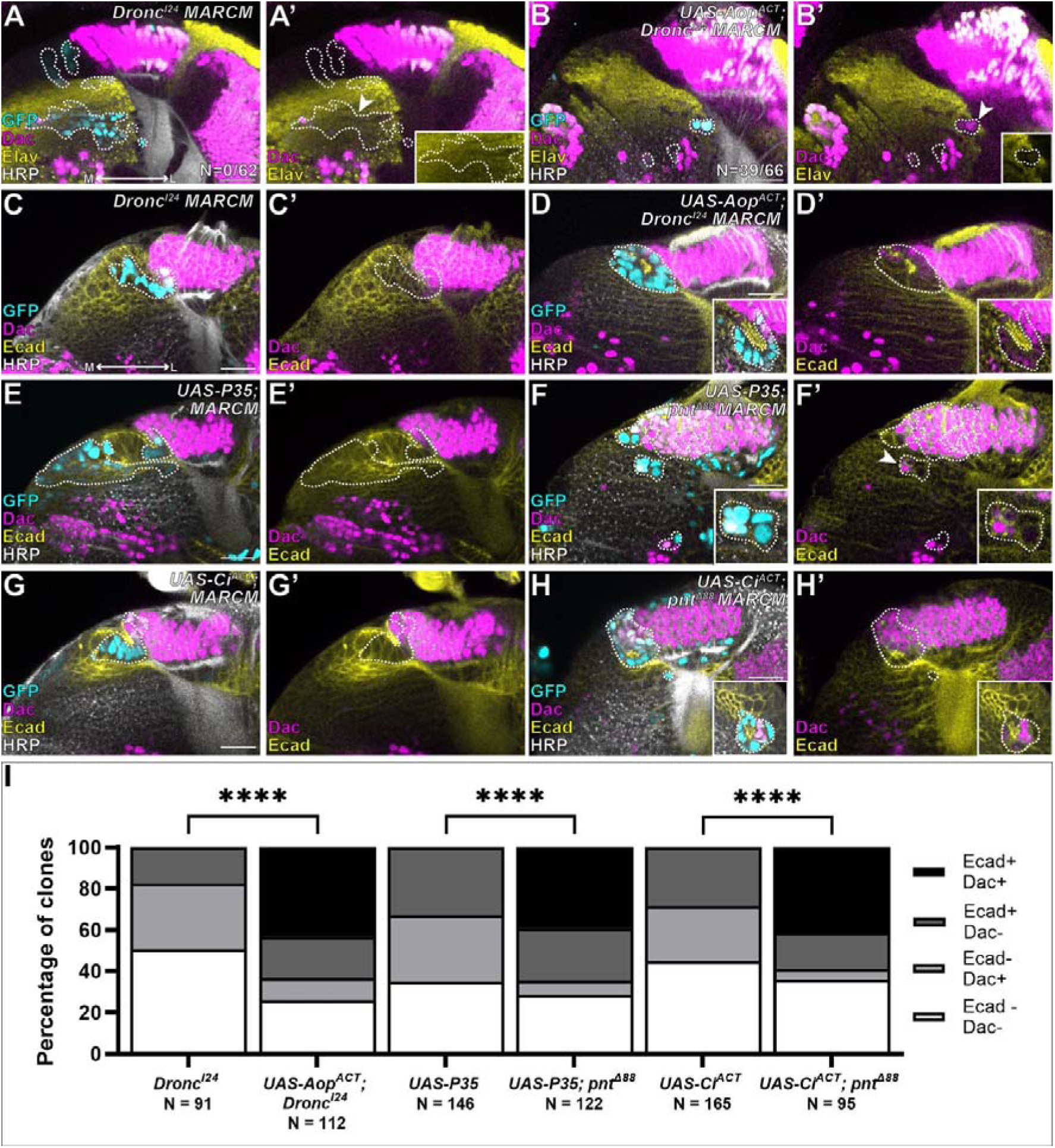
Blocking both ERK signalling and apoptosis drives ectopic expression lamina precursor differentiation. Representative cross-sections of optic lobes containing MARCM clones marked by GFP (cyan - outlined) of the following genotypes: **(A, C)** *Dronc^I24^* mutant clones **(B,D)** *Dronc^I24^* mutant clones expressing Aop^ACT^ **(E)** control clones expressing P35 **(F)** *pnt* ^Δ*88*^ mutant clones expressing P35 **(G)** control clones expressing Ci^ACT^ **(H)** *pnt* ^Δ*88*^ mutant clones expressing Ci^ACT^ stained for Dac (magenta), HRP (grey) and Elav (A,B; yellow) or Ecad (C-H; yellow). Insets in (A,B and F) show higher magnification images of indicated clones. Insets in (D,H) show clones from other focal planes of the same optic lobe which were recovered within the medulla cortex. **(I)** Quantification of the percentage of clones from (C-H) containing cells medial to the lamina furrow expressing Ecad (Ecad+ Dac-), Dac (Ecad- Dac+), co-expressing Ecad and Dac (Ecad+ Dac+) and neither Ecad nor Dac (Ecad- Dac-) (Fisher’s Exact tests, **** = P<0.0001). All optic lobes are from late L3 larvae. Scale bar = 20 μm. Medial (M), lateral (L).

Next, given our prior finding that Hh signalling promotes lamina precursor survival, we tested whether it could promote survival of cells in which ERK signalling was inhibited, thereby enabling their differentiation into ectopic lamina precursors. To do so, we generated *pnt*^Δ*88*^ clones expressing a constitutively active form of the Hh transcriptional effector Cubitus interruptus (Ci^ACT^) (Chen et al., 2000). Control clones expressing Ci^ACT^ in a wild-type background were indistinguishable from neighbouring non-clonal cells, however, *pnt*^Δ*88*^ clones expressing Ci^ACT^ contained cells ectopically co-expressing Ecad and Dac (Fig. 4G,H). We noted that the proportion of *pnt*^Δ*88*^ clones containing cells co-expressing Dac and Ecad was similar when clones expressed either P35 or Ci^ACT^ (quantified in Fig. 4I). Thus, Hh signalling is sufficient to promote survival in *pnt*^Δ*88*^ clones but does not increase the frequency of lamina-like differentiation compared to apoptosis inhibition alone.

To further validate that ERK loss-of-function clones begin differentiating into lamina precursors, we assessed the expression of additional lamina precursor markers, *glial cells missing* paralogs (*gcm* and *gcm2;* Chotard et al., 2005) by *in situ* hybridisation chain reaction (HCR). Compared to control clones expressing P35 or Ci^ACT^, *pnt*^Δ*88*^ clones expressing P35 or Ci^ACT^ showed a significant increase in the proportion of clones expressing *gcm* or *gcm2* (Fig. S3A-F), indicating that *pnt* loss-of-function clones are indeed ectopic lamina precursor cells.

Surprisingly, neither co-expression of Dac and Ecad, nor ectopic expression of *gcm*/*gcm2* was observed in clones homozygous for a second *pnt* mutant allele (*pnt*^Δ*33*^) expressing P35 or Ci^ACT^ (Fig. S3G-M). Unlike *pnt*^Δ*88*^, which disrupts all *pnt* isoforms (Brunner et al., 1994), *pnt*^Δ*33*^ deletes of an exon specific to the *pntP1* isoform (O’Neill et al., 1994). Therefore, these data indicate that ectopic lamina induction requires the disruption of all *pnt* isoforms.

Altogether, these results show that suppressing ERK signalling while promoting cell survival, either by direct inhibition of apoptosis or via Hh signalling, is sufficient to drive neuroepithelial cells to delaminate and differentiate into lamina precursors.

### ERK activity correlates with medial-lateral fate segregation in the neuroepithelium

Our data indicate that modulating ERK signalling activity is sufficient to instruct medulla versus lamina fate, with high ERK activity promoting medulla neuroblast formation and inhibition of ERK-dependent transcription results in ectopic lamina precursor formation. We therefore asked whether spatial differences in endogenous ERK activity across the neuroepithelium underlie this medial-lateral fate segregation. Using an antibody against double phosphorylated ERK (dpERK) to assess endogenous ERK activity identifies a narrow domain of ERK-active cells abutting the proneural wave (Fig. 1C) (Yasugi et al., 2010; Sato et al., 2016). However, immunohistochemistry with the dpERK may not resolve the full dynamic range of ERK signalling as cells with low levels may not be easily identifiable. We therefore employed additional reporters to better resolve variation in ERK pathway output across the neuroepithelium.

The Pnt transcription factors (PntP1, PntP2 and PntP3) are transcriptional effectors of the EGFR-ERK pathway (Wu et al., 2020). Their protein expression reflects ERK signalling output (Shwartz et al., 2013), making them a useful indirect readout of pathway activity. We therefore examined expression of GFP-tagged Pnt (Pnt::GFP), which reports expression of all Pnt isoforms (Lachance et al., 2014). In second larval instar (L2) optic lobes, Pnt::GFP was highest in the medial-most neuroepithelium and present at lower levels elsewhere (Fig. 5A,B). At early L3, Pnt::GFP remained high medially, adjacent to the youngest medulla neuroblasts (Fig. S4A, arrow), but was absent from the lateral neuroepithelium (Fig. 5C,D; outline). Notably, the lamina precursor marker Dac appeared within this Pnt::GFP-negative domain (Fig 5C’’, arrowhead). As development proceeded, the lateral neuroepithelium folded into the lamina furrow and remained Pnt::GFP negative within the trough at late L3 (Fig. 5E,F, outline), whereas delaminating lamina precursors re-expressed Pnt::GFP, consistent with a later requirement for ERK signalling in lamina neuron differentiation (Fernandes et al., 2017; Prasad et al., 2022). By ∼20 hours after puparium formation (h APF), most of the neuroepithelium had been consumed, with only a small population of Pnt::GFP-negative cells in the remnants of the furrow trough (Fig. 5G,H, outline). At this stage, Dpn expression was absent from the medulla and residual neuroepithelium (Fig. S4C), indicating that proneural wave progression had ceased before consuming the remaining neuroepithelial cells. Thus, these results suggest that ERK is active at high levels in the proneural wave, consistent with the dpERK staining, but that the rest of the neuroepithelium has some ERK activity, with the exception of cells within the trough of the lamina furrow.

**Fig. 5:**
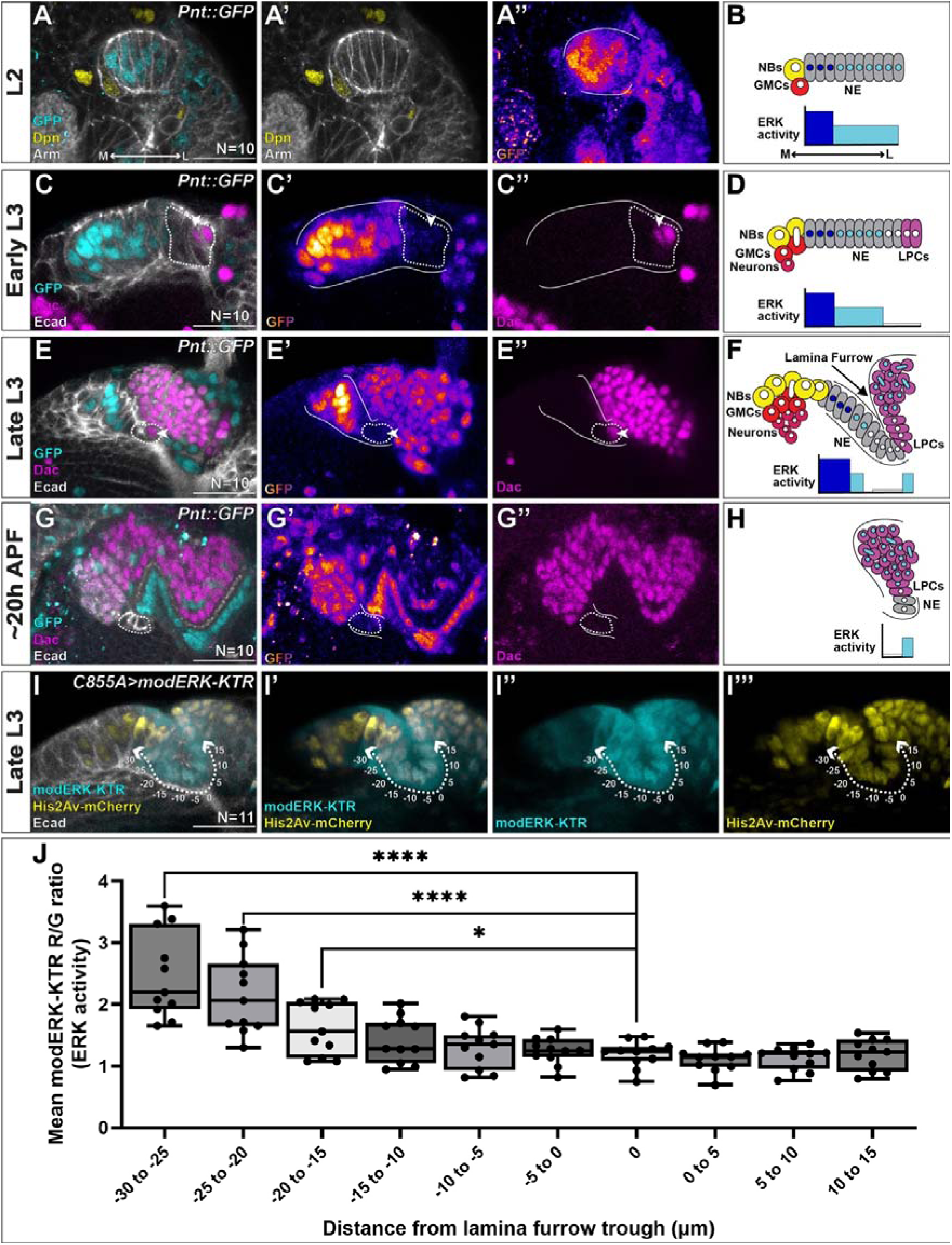
ERK signalling is excluded from the lateral neuroepithelium. **(A,C,E,G)** Representative cross-sections of optic lobes expressing Pnt::GFP (cyan, pseudo-colour) at L2 (A), early L3 (C), late L3 (E) and ∼20 h APF (G). (A) is stained for Dpn (yellow) and Arm (grey). (C,E,G) are stained for Dac (magenta) and Ecad (grey). White lines indicate the neuroepithelium. Dashed lines outline the lateral OPC lacking Pnt::GFP. Arrowheads mark the earliest Dac+ lamina precursors (N=10 lobes per timepoint). **(B,D,F,H)** Schematics of ERK activity across the OPC at the stages shown in (A,C,E,G). **(I)** Representative late L3 optic lobe expressing modERK-KTR driven by C855a-Gal4, stained for GFP (cyan), RFP (His2Av-mCherry; yellow) and Ecad (grey) (N=11). Arrows indicate positions used for quantification relative to the lamina furrow. **(J)** Quantification of modERK-KTR R/G ratio per lobe, averaged across cells within 5 μm bins up to 30 μm medial and 15 μm lateral to the lamina furrow (0 μm) (unpaired, two-tailed Mann–Whitney test, *P = 0.0387, ****P < 0.0001; whiskers = min to max). Scale bar = 20 μm. Medial (M), lateral (L), neuroblasts (NBs), ganglion mother cells (GMCs), neuroepithelial cells (NE), lamina precursor cells (LPCs).

To validate these findings and assess ERK signalling more directly, we used the modified ERK kinase translocation reporter (modERK-KTR) (Yuen et al., 2022; Wilcockson et al., 2023), in which the same promoter drives expression of a nuclear red fluorescent protein and a green fluorescent protein that shuttles between the nucleus and cytoplasm in response to ERK-dependent phosphorylation. Therefore, the nuclear red:green fluorescence ratio reports ERK activity (Yuen et al., 2022; Wilcockson et al., 2023). Driving UAS-modERK-KTR throughout the neuroepithelium using C855a-Gal4 (Manseau et al., 1997; Egger et al., 2007), we observed that the GFP signal was more cytoplasmic medial to the furrow and more nuclear within the furrow (Fig. 5I). Quantification of the nuclear Red:Green ratios across the medial-lateral axis revealed the highest ERK activity medially, then declining to the lowest levels towards the furrow trough and lateral to the trough (Fig. 5J).

Together, these data demonstrate that ERK signalling is spatially graded across the neuroepithelium, with high activity at the medial margin associated with the proneural wave, sustained suppression at the lateral margin and within the lamina furrow, and low/intermediate levels in-between. This pattern closely matches the segregation of medulla, lamina and neuroepithelial fates, and together with our observations that ERK signalling is able to induce lamina or medulla depending on whether it is inhibited or activated, supports a model in which spatial restriction of ERK activity partitions the neuroepithelium into distinct fates.

### Cortex glia secrete Argos to locally suppress ERK signalling in the neuroepithelium and promote lamina fate

Having established that ERK signalling is locally suppressed within the lamina furrow and that this suppression is required for instructing lamina fate, we next asked how ERK activity is regionally restricted within the neuroepithelium. Immature cortex glia, labelled by R54H02-Gal4 (Jenett et al., 2012; Coutinho-Budd et al., 2017), overlie the neuroepithelium and, at L3, extend a long process deep into the lamina furrow (Fig. 6A). These glia express the secreted EGF antagonist Argos (Aos; visualised with *aos-lacZ;* Fig. 6B) (Morante et al., 2013), raising the possibility that they locally suppress EGFR-ERK signalling within the furrow.

**Fig. 6:**
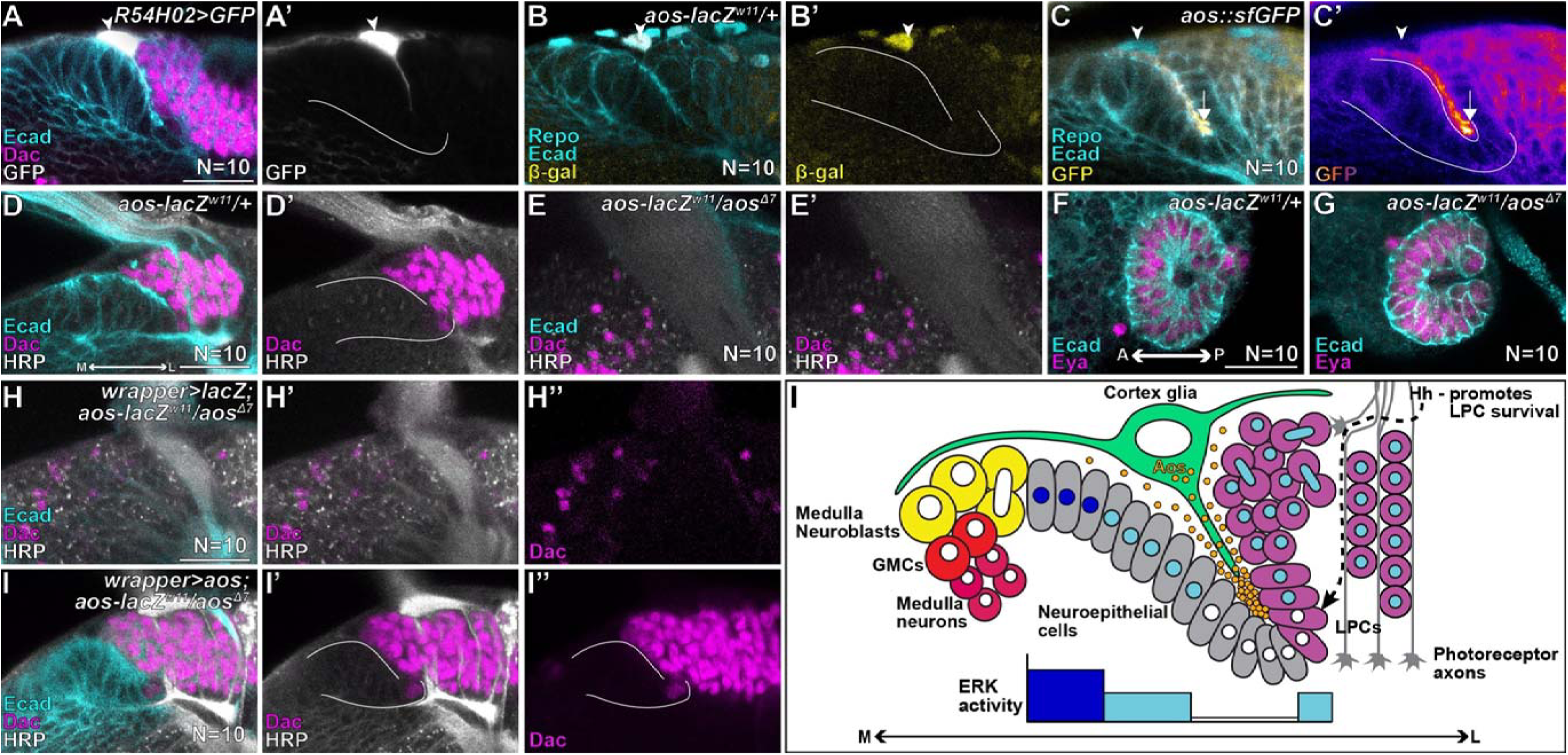
Restoring *aos* in cortex glia in an *aos* mutant rescues lamina development. **(A)** Representative late L3 optic lobe with R54H02-Gal4 driving expression of GFP (grey) in cortex glia. **(B,C)** Representative optic lobes from late L3 *aos-lacZ^w11^* heterozygous larvae (B) and *aos::sfGFP* larvae stained for Repo (cyan - nuclear), Ecad (cyan – apico-lateral membranes) and β-galactosidase (B; yellow) or GFP (C; yellow, C’, pseudo-colour). White arrowheads mark cortex glia in (A,B,C). **(D,E)** Representative optic lobes from late L3 *aos-lacZ^w11^* mutant heterozygous larvae (D) and *aos-lacZ^w11^/aos* ^Δ*7*^ mutant transheterozygous larvae (E) stained for Ecad (cyan) Dac (magenta) and HRP (grey). **(F,G)** Representative optic lobes from L2 *aos-lacZ^w11^* heterozygous larvae (F; N=10) and *aos-lacZ^w11^/aos* ^Δ*7*^ larvae (G; N=10) stained for Ecad (cyan) and the OPC neuroepithelium marker Eya (magenta). **(H,I)** Representative late L3 *aos-lacZ^w11^/aos* ^Δ*7*^ optic lobes with wrapper-Gal4 driving expression of *lacZ* (H) or *aos* (I) stained for Ecad (cyan), Dac (magenta) and HRP (grey). White lines indicate the neuroepithelium up to the lateral lamina furrow. **(I)** Schematic depicting our proposed model for lamina lineage specification where cortex glia secrete Aos to suppress EGFR-ERK signalling locally in the lamina furrow. This in turn instructs lamina precursor development but necessitates photoreceptor-derived Hh to promote LPC survival. All images are transverse cross-sections. N=10 lobes for each condition, scale bar = 20 μm. Medial (M), lateral (L), neuroblasts (NBs), ganglion mother cells (GMCs), lamina precursor cells (LPCs).

To test this, we examined Aos protein localisation using a protein trap encoded by *aos::sfGFP* (Sarov et al., 2016). Aos::sfGFP accumulated at high levels within the trough of the lamina furrow, forming a gradient peaking at the site where ERK activity is absent (Fig. 6C). This distribution is suggestive of local secretion by the cortex glia and accumulation of Aos within the furrow. In line with the hypothesis that Aos locally represses ERK signalling to enable lamina specification, trans-heterozygous *aos* mutants (*aos-lacZ^w11^/aos* ^Δ*7*^; (Freeman et al., 1992) failed to develop a lamina at L3, compared to heterozygous controls (*aos-lacZ^w11^/+*), as reported previously (Fig. 6D,E) (Yasugi et al., 2010; Morante et al., 2013). Importantly, this defect did not reflect an earlier failure in neuroepithelial development, as the neuroepithelial crescent was present and indistinguishable from controls at earlier stages (Fig. 6F,G). These data indicate that *aos* is specifically required during L3 for lamina formation.

Since *aos* is required for lamina precursors to be specified, we asked whether modulating Aos levels would affect the relative sizes of the lamina and medulla. In particular, we tested whether increasing Aos levels in cortex glia is sufficient to bias neuroepithelial fate allocation towards the lamina at the expense of the medulla. To do so, we overexpressed *aos* in cortex glia using R54H02-Gal4. Compared to control optic lobes, overexpression of *aos* in cortex glia led to a significant expansion of the lamina together with a corresponding reduction in medulla size (Fig. S5A,B; quantified in C,D). Thus, increasing Aos levels in cortex glia is sufficient to bias neuroepithelial fate towards lamina at the expense of medulla.

Although cortex glia are uniquely positioned to deliver Aos directly to the lamina furrow, they are not the only cells which express *aos* at L3. In addition to cortex glia, *aos-lacZ* expression was detected in differentiating lamina neurons and surface glia (Fig. 6B)(Prasad et al., 2022). To determine whether cortex glia represent the relevant source of Aos, we restored *aos* expression specifically in cortex glia using wrapper-Gal4 (Lee et al., 2018) in an *aos* mutant background (*aos-lacZ^w11^/aos* ^Δ*7*^), which otherwise lacks the lamina. Remarkably, expressing *aos* in cortex glia alone fully rescued lamina development (Fig. 6H-I).

Together, these results demonstrate that cortex glia secrete Aos to locally suppress EGFR-ERK signalling within the lamina furrow, thereby partitioning the neuroepithelium into lamina and medulla domains.

## Discussion

In this study, we address how a single neuroepithelium is partitioned to generate two distinct visual processing centres, the lamina and medulla. Previous work has proposed a model in which different spatially restricted signals instruct these fates on each side of the epithelium, with EGFR-ERK signalling promoting medulla development on the medial side (Yasugi et al., 2008; Egger et al., 2010; Yasugi et al., 2010; Sato et al., 2016; Jörg et al., 2019) and photoreceptor-derived Hedgehog (Hh) inducing lamina formation on the lateral side (Huang and Kunes, 1996; Huang and Kunes, 1998). Here, we show that Hh signalling is not instructive for lamina precursor specification or cell cycle progression, but instead acts permissively by promoting survival of cells already specified as lamina precursors from the lateral neuroepithelium. In contrast, we find that EGFR-ERK signalling, which drives proneural wave progression and medulla neuroblast formation, must be suppressed to enable neuroepithelial cells to differentiate into lamina precursors and that suppression of ERK together with promotion of cell survival is sufficient to specify lamina fate. Finally, we demonstrate that this suppression is mediated by cortex glia, which secrete the EGF antagonist Aos to locally inhibit EGFR-ERK signalling locally in the lateral neuroepithelium. Thus, glia extrinsically pattern the neuroepithelium by spatially restricting ERK signalling, enabling a single signalling pathway to instruct the formation of distinct visual processing centres.

### Spatially modulating ERK activity instructs distinct fates in the neuroepithelium

Several prior studies have shown that high EGFR-ERK activity drives medulla neuroblast formation (Yasugi et al., 2008; Egger et al., 2010; Yasugi et al., 2010; Sato et al., 2016; Jörg et al., 2019). Here we show that complete suppression of ERK activity instructs lamina precursor development, whereas partial suppression of ERK activity maintains neuroepithelial fate (Fig. S3G-J) (Weng et al., 2012). Thus, rather than acting as a binary switch, ERK signalling functions as a graded instructive cue whose spatial modulation partitions the neuroepithelium into discrete domains. These findings raise the question of how different levels of ERK activity are interpreted at the transcriptional level. Our observation that complete loss of all *pnt* isoforms (*pnt*^Δ*88*^), but not loss of *pntP1* alone (*pnt*^Δ*33*^), is sufficient to induce ectopic lamina differentiation suggests that multiple Pnt isoforms cooperate to repress lamina fate.

### Glia as extrinsic regulators of neuroepithelial patterning

Glia are increasingly recognised as active regulators of neural development (Lago-Baldaia et al., 2020). The Drosophila lamina is a striking example of this, as multiple glial populations coordinate and drive neuronal differentiation (Fernandes et al., 2017; Prasad et al., 2022; Donoghue et al., 2025). Our findings extend this paradigm by showing that cortex glia act earlier to partition the neuroepithelium itself, by spatially restricting ERK signalling through Aos secretion.

Cortex glia have previously been implicated in regulating neuroepithelial proliferation and the timing of medulla neuroblast formation through secretion of EGF (Morante et al., 2013). Here, we show that they also inhibit EGFR-ERK signalling locally to enable lamina precursor differentiation, highlighting a dual role in both promoting and restricting pathway activity depending on context. Similar functions in regulating neural stem cell proliferation and differentiation have been shown for microglia in vertebrate nervous system development (Morgan et al., 2004; Huang et al., 2012).

An unresolved question is how cortex glia restrict EGFR-ERK activity to the lateral neuroepithelium, given that they extend broadly over the entire neuroepithelial surface. We observe that Aos protein accumulates at high levels within the lamina furrow (Fig. 6C). One possibility is that Aos is actively trafficked along glial processes that extend into the furrow. Alternatively, the architecture of the furrow itself, where apical surfaces of neuroepithelial cells are closely apposed, may more efficiently capture and concentrate secreted Aos.

### Photoreceptor-derived Hh promotes lamina precursor survival in the absence of ERK activity

Our data demonstrate the Hh signalling is not instructive for specifying lamina precursor fate but instead that it acts permissively by promoting survival of cells that have already been specified as lamina precursors. Nonetheless, the reduced levels of Dac observed in lamina precursors in the absence of Hh signalling suggest that it may contribute to stabilising or reinforcing lamina identity once initiated.

Our findings are consistent with known roles of Hh signalling in inhibiting apoptosis in other contexts, including the Drosophila wing disc and in vertebrate neuroepithelia (Thibert et al., 2003; Cayuso et al., 2006; Lu et al., 2017). In the wing pouch Hh signalling cell-autonomously promotes expression of anti-apoptotic factor *Death-associated inhibitor of apoptosis 1* (*Diap1)* while also repressing expression of the pro-apoptotic gene *head involution defective (hid)* (Lu et al., 2017. In parallel, ERK signalling has also been shown to promote cell survival in multiple contexts by regulating Hid (Bergmann et al., 1998; Domínguez et al., 1998; Kurada and White, 1998; Bergmann et al., 2002; Brown and Freeman, 2003; Crossman et al., 2018). Our findings suggest that, within the lamina furrow suppression of ERK activity renders neuroepithelial cells and differentiating lamina precursors vulnerable to apoptosis, necessitating an alternative survival signal. Hh signalling, activated locally at the lamina furrow by photoreceptor axons, fulfils this role.

Interestingly, lamina precursors at the tips of the lamina do not require Hh signalling for survival (Fig. S1B-D), indicating regional differences in survival requirements. This suggests that an alternative survival signal operates at the lamina tips. Given the proximity of these regions to domains of *decapentaplegic* (*dpp*) expression in the neuroepithelium (Yoshida et al., 2005; Suzuki et al., 2016b), it is possible that Dpp provides a compensatory survival signal in this context as it does in the wing pouch (Bryant, 1988; Adachi-Yamada et al., 1999).

Together, these findings reveal a division of labour between signalling pathways: ERK activity instructs fate, while Hh signalling ensures survival of appropriately specified cells. This coordination ensures that lamina precursors are generated at the correct location and maintained by the arrival of photoreceptor axons.

## Materials and methods

### Drosophila stocks and maintenance

*D. melanogaster* strains and crosses were reared on standard cornmeal medium and reared at 25°C unless otherwise noted. The following mutant and transgenic flies were used in combination or recombined in this study; ({ } enclose individual genotypes separated by commas).

Canton S*, {y,w,hsflp^122^; sp/Cyo; TM2/TM6B}, {y,w; sp/Cyo, Bacc-GFP; Dr/TM6C},* (from BDSC: 36349), {*;Ecad::GFP;*} (a gift from Y. Mao), {*; ptc::GFP/Cyo; hh^ts2^/TM6C*} (a gift from G. Struhl), {*w*; pnt-GFP;*} (*Pnt::GFP*, BSDC: 42680), *{w; UAS-Aop^ACT^;*} (BDSC: 5789), *{y^1^,sc*,v^1^,sev^21^;; UAS-aop^RNAi^*} (BDSC: 34909), {*w*;;UAS-EGFR* ^λ*top*^} (EGFR^ACT^, BDSC: 59843), {*y,w,hsflp^122^, Tub>Gal4, UAS-nls.GFP; Tub>Gal80, FRT40A*}, {*y,w,hsflp^122^, Tub>Gal4, UAS-mCherry.nls; Tub>Gal80, FRT40A*}, {*y,w,hsflp^122^, UAS-nls.RFP; Tub>Gal80, FRT42D; Tub>Gal4*}, {*y,w,hsflp^122^, Tub>Gal4, UAS-nls.GFP;; Tub>Gal80, FRT80B/TM2*}, {*y,w,hsflp^122^, Tub>Gal4, UAS-nls.GFP;; Tub>Gal80, FRT82B/TM2*} (gifts from M. Amoyel), {;*smo^3^, FRT40A/CyO;*} (a gift from C. Boekel), {*y,w*;;*Dronc^I24^, FRT80B/6B*} (a gift from M. Amoyel), {;*aop^EX18^, FRT40A/CyO;*} (a gift from J. Treisman), {*w*;FRT40A;*} (a gift from M. Amoyel), {*y^1^; FRT42D, ptc^S2^/CyO*} (BDSC: 6332), {*;;pnt*^Δ*88*^/*TM3*} (BDSC: 861), {*w*;; FRT82B, pnt*^Δ*33*^*/TM6B*} (a gift from M. Amoyel), {*y,w,hsflp^122^; sp/CyO; FRT82B/TM6B*} (a gift from M. Amoyel), {*w^1118^;; R54H02-Gal4*} (cortex glia; BDSC: 45784), {*w*; wrapper-Gal4/CyO*} (BSDC: 93483), {*w*;UAS-p35;*} (BSDB: 5072), {*w*;;UAS-P35*} (BSDC: 5073), {*w*; UAS-ci^m1-3*103^*} (ci^ACT^, BDSC: 32571), {*w^1118^;;C855a-Gal4*} (BSDC: 6990),{*y^1^,w*; UAS-modERK-KTR-T2A-His2AV-mCh/CyO;*} (modERK-KTR, BDSC: 95286), {*y,w,hsflp^122^*; *sp/CyO*; *pxb-T2AGal4/TM6B*} (a gift from C. Desplan), {*y,w,hsflp^122^*; *Tub-Gal80^ts^/CyO*; *TM2/TM6B*} (a gift from M. Amoyel), {*;UAS-nls.LacZ;*} (BSDC:3955), {*;;UAS-nls.LacZ*} (BSDC:3956), {*y^1^,w*; UAS-aos; UAS-aos*} (BSDC: 5363), {*w^1118^; UAS-GFP.nls*} (BSDC: 4775), {*w*; 10XUAS-CD8::GFP;*} (BSDC: 32186), {;; *aos::sfGFP*} (Vienna Drosophila Resource Centre [VDRC]: 318382), {*w;;aos-LacZ^w11^/TM3*} (BSDC: 2513), {*;;aos* ^Δ*7*^*/TM3*} (BSDC: 1004), {*;;aop::sfGFP*} (VDRC: 318050).

Crosses using *pxb-Gal4^ts^* (Fig. 3) were incubated at 18°C for 7 days after egg laying (AEL), at which point they were shifted to 29°C for 48 hours to relieve inhibition of GAL4 by GAL80 and restrict *pxb-Gal4* expression to L3 stage.

For full experimental genotypes and conditions see Table S1.

### Timed collections

*Pnt::GFP* flies were flipped onto fresh food and allowed to lay for 3 hours. Larvae from these collections were dissected and fixed 72 hours later (second larval instar; L2), 96 hours later (Early L3), 120 hours later (Late L3) and at 20 hours after puparium formation (APF).

### Clonal analysis

MARCM clones were induced by heat-shocking larvae at 37°C either 3 days AEL for 60 minutes ({*FRT40A aop^XE18^*}, {*FRT80B Dronc^I24^*}, {*UAS-aop^ACT^; FRT80B Dronc^I24^*}, {*UAS-P35; FRT82B*},{*UAS-ci^ACT^; FRT82B*}, {*UAS-P35*; *FRT82B pnt*^Δ*88*^}, {*UAS-ci^ACT^*; *FRT82B pnt*^Δ*88*^}, {*UAS-P35*; *FRT82B pnt*^Δ*33*^}, {*UAS-ci^ACT^*; *FRT82B pnt*^Δ*33*^}) or 2 days AEL for 90 minutes ({*FRT40A smo^3^*}, {*FRT40A smo^3^*; *UAS-P35*}, {*FRT42D ptc^S2^*}). All MARCM crosses were raised at 25°C until dissection.

### Immunohistochemistry, antibodies and microscopy

We dissected eye-optic lobe complexes in 1× phosphate-buffered saline (PBS), fixed in 4% formaldehyde for 20 min, blocked in 5% normal donkey serum, and incubated in primary antibodies diluted in block for two nights at 4°C. Samples were then washed in 1× PBS with 0.5% Triton-X (PBSTx), incubated in secondary antibodies diluted in block, washed in PBSTx and mounted in SlowFade (Life Technologies).

When performing phospho-ERK stains, dissections were performed in a phosphatase inhibitor buffer as detailed in Amoyel et al. (2016)

The following antibodies were obtained from the Developmental Studies Hybridoma Bank (DHSB), created by the NICHD of the NIH and maintained at The University of Iowa, Department of Biology, Iowa City, IA 52242: mouse anti-Arm (1:200, N2 7A1 Armadillo, Wieschaus, E.), mouse anti-Dac^2-3^ (1:100, mAbdac2-3, Rubin, G.M.), rat anti-Ecad (1:20, DCAD2, Uemura, T.), rat anti-Elav (1:100, Rat-Elav-7E8A10 anti-elav, Rubin, G.M), mouse anti-Eya (1:100, eya10H6, Benzer, S. / Bonini, N.M), mouse-anti-GFP (1:500, DSHB-GFP-4C9, DSHB).

In addition, we also used the following primary antibodies in this study: mouse anti-β-galactosidase (1:125; Promega #Z3781), guinea pig anti-Dac (1:1000, a gift from C. Desplan), rabbit anti-Dcp-1 (1:100; Cell Signalling #9578), rat anti-Dpn (1:250, abcam ab195172), rabbit-anti-GFP (1:500; Thermo Fisher Scientific #A6455), rat anti-Repo (1:500, a gift from C. Desplan), chicken anti-RFP (1:500; Rockland #600-901-379s), rabbit anti-p-H3 (ser10) (1:100, Cell signalling #9701), rabbit anti-Phospho-p44/42 MAPK (Erk1/2) (Thr202/Tyr204), (1:100, Cell Signaling #9101), AlexaFluor405 conjugated Goat Anti-HRP (1:100, Jackson Immunolabs).

We used the following secondary antibodies at a concentration of 1:500 in this study: Alexa Fluor® 488 AffiniPure Donkey Anti-Rabbit IgG (H+L) (Jackson ImmunoResearch, #711-545-152), Rhodamine Red™-X (RRX) AffiniPure® Donkey Anti-Rabbit IgG (H+L) (Jackson ImmunoResearch, #711-295-152), Rhodamine Red™-X (RRX) AffiniPure® Donkey Anti-Guinea Pig IgG (H+L) (Jackson ImmunoResearch, #706-295-148), Donkey anti-Mouse IgG (H+L) Highly Cross-Adsorbed Secondary Antibody, Alexa Fluor™ 488 (Invitrogen, #A21202), Alexa Fluor® 647 AffiniPure® Donkey Anti-Rat IgG (H+L) (Jackson ImmunoResearch, #712-605-153), Donkey anti-Mouse IgG (H+L) Highly Cross-Adsorbed Secondary Antibody, Alexa Fluor™ 568 (Invitrogen, #A10037), Donkey anti-Mouse IgG (H+L) Highly Cross-Adsorbed Secondary Antibody, Alexa Fluor™ 647 (Invitrogen, #A31571), Goat anti-Chicken IgY (H+L) Secondary Antibody, Alexa Fluor™ 568 (Invitrogen, #A11041). Images were acquired using a Zeiss 800 confocal microscope with a 40× objective.

### EdU incorporation Assay

L3 stage larvae were dissected in fresh Schneider’s Medium (Merck - S0146). Dissected brains were incubated in a solution of EdU (10μM - Abcam - ab146186) dissolved in Schneider’s medium for 30 minutes at room temperature with shaking. The immunostaining procedure outlined above was then carried out until the 1X PBS wash. The Click reaction was initiated just before washing in 1X PBS. Brains were incubated in Click solution (2.5μM picolyl Azide 405nm [DC Bioscience –1308-1], 0.1mM Tris[benzyltriazolylmethyl]amine, 2mM Sodium Ascorbate, and 1mM Copper Sulphate in 1X PBS) for 30 minutes at room temperature with shaking. The brains were further washed in 1X PBS for 30 minutes.

### *In situ* HCR

To assess ectopic expression of additional LPC markers in MARCM clones (Fig. S3) we designed HCR probes against *gcm* and *gcm2*. We designed 20 antisense probe pairs (see supplementary table 2) against each target gene, tiled along the annotated transcripts but excluding regions of strong sequence similarity to other transcripts, with the corresponding initiator sequences for amplifiers B3 and B5 (Choi et al., 2018). We purchased HCR probes as DNA oligos from Thermo Fisher (at 100 μM in water and frozen). All probe sequences are included in supplementary file 1.

Eye-optic lobe complexes were dissected, fixed, and permeabilised as above. Samples were incubated in probe hybridisation buffer at 37°C for 30 min before being incubated with probes at 0.01 µM at 37°C overnight. Samples were washed four times for 15 min at 37°C with probe wash buffer and then two times for 5 min with 5× saline-sodium citrate with 0.001% Tween 20 solution (to make a 20× SSCT solution in distilled H_2_O, 58.44 g/mol sodium chloride, 294.10 g/mol sodium citrate, pH adjusted to 7 with 14 N hydrochloric acid, with 0.001% Tween 20). Samples were incubated in an amplification buffer for 10 min.

Hairpins H1 and H2 for each probe were snap-cooled (hairpins were heated to 95°C for 90 s and cooled to room temperature for 30 min) separately to avoid hairpin oligomerisation. About 12 pmol of each hairpin was added to samples in an amplification buffer and incubated overnight at room temperature. Samples were washed for 10 min in SSCT and then incubated in darkness at room temperature with 1:500 dilution of DAPI (Sigma: D9542) for 45 min. Samples were washed in 1× PBS for 30 min and mounted as above.

### Quantification and statistical analyses

We used Fiji-ImageJ (Schindelin et al., 2012) to process and quantify confocal images as described below. We used Adobe Photoshop and Adobe Illustrator software to prepare figures. We used GraphPad Prism8 software to perform statistical tests. In all graphs, whiskers indicate the minimum and maximum values. N values are indicated on graphs.

### Onset of Dac expression

To quantify the onset of Dac expression in the lamina furrow, we used the cell at the trough of the lamina furrow as a starting point (cell position 0) and assessed the onset of Dac expression up to four cells medially (positions −1 to −4) and up to three cells laterally (positions 1 to 3).

### MARCM clone location quantification

To quantify the position of control clones, *smo^3^* clones, and *smo^3^* clones expressing UAS-P35, we used the tips of the lamina as a starting point and measured the location of the centre of each clone along the anterior circumference up to the centre of the lamina. We then took the proportionate distance along the lamina circumference such that the scale along the lamina circumference became 0.0 at the lamina tips and 1.0 at the centre of the lamina.

### Ectopic LPC quantification

For quantification of LPC marker expression in MARCM clones (Fig. 4, Fig. S3) clones within the OPC medial to the lamina furrow and medulla cortex were counted and individually assessed to determine whether they contained cells expressing each marker.

### Quantifications of modERK-KTR R/G ratio

modERK-KTR R/G ratios were measured in fixed optic lobes from late L3 larvae by manually segmenting nuclei on ImageJ using the freehand selection tool at the plane with the largest nuclear diameter. Nuclei were measured up to 30 µm medial and 15 µm lateral to the trough of the lamina furrow. The distance between nuclei along the lamina furrow and the nucleus at the trough of the lamina furrow on the same plane was measured using the line tool.

Nuclei medial to the trough were converted to negative values. R/G ratios were averaged across 5 µm bins to produce mean R/G ratios for group.

### Quantifications of neuropil area

Medulla and lamina area were measured using the freehand selection tool on ImageJ. Area of each neuropil was measured across 5 focal planes spanning 15 µm in nine optic lobes for each condition. In all samples, area measurements were consistently made within the anterior portion of the OPC containing the central region of the lamina crescent.

## Acknowledgements

We are grateful to M. Amoyel, R. Poole and members of the Fernandes, Amoyel and Poole labs for critical feedback on the work. Stocks obtained from the Bloomington Drosophila Stock Center (NIH P40OD018537) were used in this study. Monoclonal antibodies obtained from the Developmental Studies Hybridoma Bank, created by the NICHD of the NIH and maintained at The University of Iowa, were used in this study.

## References

Adachi-Yamada, T., Fujimura-Kamada, K., Nishida, Y. and Matsumoto, K. (1999) ‘Distortion of proximodistal information causes JNK-dependent apoptosis in Drosophila wing’, Nature 400(6740): 166–9.

Amoyel, M., Anderson, J., Suisse, A., Glasner, J. and Bach, E. A. (2016) ‘Socs36E Controls Niche Competition by Repressing MAPK Signaling in the Drosophila Testis’, PLoS Genet 12(1): e1005815.

Apitz, H. and Salecker, I. (2014) ‘A challenge of numbers and diversity: neurogenesis in the Drosophila optic lobe’, J Neurogenet 28(3-4): 233–49.

Bergmann, A., Agapite, J., McCall, K. and Steller, H. (1998) ‘The Drosophila gene hid is a direct molecular target of Ras-dependent survival signaling’, Cell 95(3): 331–41.

Bergmann, A., Tugentman, M., Shilo, B. Z. and Steller, H. (2002) ‘Regulation of cell number by MAPK-dependent control of apoptosis: a mechanism for trophic survival signaling’, Dev Cell 2(2): 159–70.

Bertet, C., Li, X., Erclik, T., Cavey, M., Wells, B. and Desplan, C. (2014) ‘Temporal patterning of neuroblasts controls Notch-mediated cell survival through regulation of Hid or Reaper’, Cell 158(5): 1173–1186.

Bostock, M. P., Prasad, A. R., Donoghue, A. and Fernandes, V. M. (2022) ‘Photoreceptors generate neuronal diversity in their target field through a Hedgehog morphogen gradient in Drosophila’, Elife 11.

Brown, Katherine E and Freeman, Matthew (2003) ‘Egfr signalling defines a protective function for ommatidial orientation in the Drosophila eye’.

Brunner, Damian, Dücker, Klaus, Oellers, Nadja, Hafen, Ernst, Scholzi, Henrike and Klambt, Christian (1994) ‘The ETS domain protein pointed-P2 is a target of MAP kinase in the sevenless signal transduction pathway’, Nature 370(6488): 386–389.

Bryant, P. J. (1988) ‘Localized cell death caused by mutations in a Drosophila gene coding for a transforming growth factor-beta homolog’, Dev Biol 128(2): 386–95.

Cayuso, Jordi, Ulloa, Fausto, Cox, Barny, Briscoe, James and Martí, Elisa (2006) ‘The Sonic hedgehog pathway independently controls the patterning, proliferation and survival of neuroepithelial cells by regulating Gli activity’.

Chen, Y. and Struhl, G. (1996) ‘Dual roles for patched in sequestering and transducing Hedgehog’, Cell 87(3): 553–63.

Chen, Yang, Goodman, RH and Smolik, Sarah M (2000) ‘Cubitus interruptus requires Drosophila CREB-binding protein to activate wingless expression in the Drosophila embryo’, Molecular and cellular biology 20(5): 1616–1625.

Chen, Yu and Struhl, Gary (1998) ‘In vivo evidence that Patched and Smoothened constitute distinct binding and transducing components of a Hedgehog receptor complex’, Development 125(24): 4943–4948.

Choi, H. M. T., Schwarzkopf, M., Fornace, M. E., Acharya, A., Artavanis, G., Stegmaier, J., Cunha, A. and Pierce, N. A. (2018) ‘Third-generation in situ hybridization chain reaction: multiplexed, quantitative, sensitive, versatile, robust’, Development 145(12).

Chotard, Carole, Leung, Wendy and Salecker, Iris (2005) ‘glial cells missing and gcm2 cell autonomously regulate both glial and neuronal development in the visual system of Drosophila’, Neuron 48(2): 237–251.

Cohen, M., Briscoe, J. and Blassberg, R. (2013) ‘Morphogen interpretation: the transcriptional logic of neural tube patterning’, Curr Opin Genet Dev 23(4): 423–8.

Courgeon, M. and Desplan, C. (2019) ‘Coordination between stochastic and deterministic specification in the Drosophila visual system’, Science 366(6463).

Coutinho-Budd, Jaeda C, Sheehan, Amy E and Freeman, Marc R (2017) ‘The secreted neurotrophin Spätzle 3 promotes glial morphogenesis and supports neuronal survival and function’, Genes & development 31(20): 2023–2038.

Crossman, Samuel H, Streichan, Sebastian J and Vincent, Jean-Paul (2018) ‘EGFR signaling coordinates patterning with cell survival during Drosophila epidermal development’, PLoS biology 16(10): e3000027.

Domínguez, María, Wasserman, Jonathan D and Freeman, Matthew (1998) ‘Multiple functions of the EGF receptor in Drosophila eye development’, Current biology 8(19): 1039–1048.

Donoghue, Alicia, Mosby, Lewis S, Lago-Baldaia, Inês, Hodgetts, Tamara, Ursu, Evelina, Erten, Zeynep, Hadjivasiliou, Zena and Fernandes, Vilaiwan M (2025) ‘Morphogen and juxtacrine signalling dynamically integrate to specify cell fates with single-cell resolution’, bioRxiv: 2025.09. 15.676088.

Egger, B., Gold, K. S. and Brand, A. H. (2010) ‘Notch regulates the switch from symmetric to asymmetric neural stem cell division in the Drosophila optic lobe’, Development 137(18): 2981–7.

Egger, Boris, Boone, Jason Q, Stevens, Naomi R, Brand, Andrea H and Doe, Chris Q (2007) ‘Regulation of spindle orientation and neural stem cell fate in the Drosophila optic lobe’, Neural development 2(1): 1.

Erclik, T., Li, X., Courgeon, M., Bertet, C., Chen, Z., Baumert, R., Ng, J., Koo, C., Arain, U., Behnia, R. et al. (2017) ‘Integration of temporal and spatial patterning generates neural diversity’, Nature 541(7637): 365–370.

Fernandes, V. M., Chen, Z., Rossi, A. M., Zipfel, J. and Desplan, C. (2017) ‘Glia relay differentiation cues to coordinate neuronal development in Drosophila’, Science 357(6354): 886–891.

Fischbach, K-F and Dittrich, APM (1989) ‘The optic lobe of Drosophila melanogaster. I. A Golgi analysis of wild-type structure’, Cell and tissue research 258(3): 441–475.

Freeman, Matthew, Klämbt, Christian, Goodman, Corey S and Rubin, Gerald M (1992) ‘The argos gene encodes a diffusible factor that regulates cell fate decisions in the Drosophila eye’, Cell 69(6): 963–975.

Huang, T., Cui, J., Li, L., Hitchcock, P. F. and Li, Y. (2012) ‘The role of microglia in the neurogenesis of zebrafish retina’, Biochem Biophys Res Commun 421(2): 214–20.

Huang, Zhen and Kunes, Samuel (1996) ‘Hedgehog, transmitted along retinal axons, triggers neurogenesis in the developing visual centers of the Drosophila brain’, Cell 86(3): 411–422.

Huang, Zhen and Kunes, Samuel (1998) ‘Signals transmitted along retinal axons in Drosophila: Hedgehog signal reception and the cell circuitry of lamina cartridge assembly’, Development 125(19): 3753–3764.

Jenett, Arnim, Rubin, Gerald M, Ngo, Teri-TB, Shepherd, David, Murphy, Christine, Dionne, Heather, Pfeiffer, Barret D, Cavallaro, Amanda, Hall, Donald and Jeter, Jennifer (2012) ‘A GAL4-driver line resource for Drosophila neurobiology’, Cell reports 2(4): 991–1001.

Jörg, David J, Caygill, Elizabeth E, Hakes, Anna E, Contreras, Esteban G, Brand, Andrea H and Simons, Benjamin D (2019) ‘The proneural wave in the Drosophila optic lobe is driven by an excitable reaction-diffusion mechanism’, Elife 8: e40919.

Karim, Felix D, Chang, Henry C, Therrien, Marc, Wassarman, David A, Laverty, Todd and Rubin, Gerald M (1996) ’A screen for genes that function downstream of Ras1 during Drosophila eye development’, Genetics 143(1): 315–329.

Konstantinides, N., Holguera, I., Rossi, A. M., Escobar, A., Dudragne, L., Chen, Y. C., Tran, T. N., Martinez Jaimes, A. M., Ozel, M. N., Simon, F. et al. (2022) ‘A complete temporal transcription factor series in the fly visual system’, Nature 604(7905): 316–322.

Kurada, P. and White, K. (1998) ‘Ras promotes cell survival in Drosophila by downregulating hid expression’, Cell 95(3): 319–29.

Lachance, Jean-François Boisclair, Peláez, Nicolás, Cassidy, Justin J, Webber, Jemma L, Rebay, Ilaria and Carthew, Richard W (2014) ‘A comparative study of Pointed and Yan expression reveals new complexity to the transcriptional networks downstream of receptor tyrosine kinase signaling’, Developmental Biology 385(2): 263–278.

Lago-Baldaia, I., Fernandes, V. M. and Ackerman, S. D. (2020) ‘More Than Mortar: Glia as Architects of Nervous System Development and Disease’, Front Cell Dev Biol 8: 611269.

Lee, Pei-Tseng, Zirin, Jonathan, Kanca, Oguz, Lin, Wen-Wen, Schulze, Karen L, Li-Kroeger, David, Tao, Rong, Devereaux, Colby, Hu, Yanhui and Chung, Verena (2018) ‘A gene-specific T2A-GAL4 library for Drosophila’, Elife 7: e35574.

Lee, Tzumin and Luo, Liqun (2001) ‘Mosaic analysis with a repressible cell marker (MARCM) for Drosophila neural development’, Trends in neurosciences 24(5): 251–254.

Li, Xin, Erclik, Ted, Bertet, Claire, Chen, Zhenqing, Voutev, Roumen, Venkatesh, Srinidhi, Morante, Javier, Celik, Arzu and Desplan, Claude (2013) ‘Temporal patterning of Drosophila medulla neuroblasts controls neural fates’, Nature 498(7455): 456–462.

Lu, Juan, Wang, Dan and Shen, Jie (2017) ‘Hedgehog signalling is required for cell survival in Drosophila wing pouch cells’, Scientific Reports 7(1): 11317.

Manseau, Lynn, Baradaran, Ali, Brower, Danny, Budhu, Anuradha, Elefant, Felice, Phan, Huy, Philp, Alastair Valentine, Yang, Mingyao, Glover, David and Kaiser, Kim (1997) ‘GAL4 enhancer traps expressed in the embryo, larval brain, imaginal discs, and ovary of Drosophila’, Developmental dynamics: an official publication of the American Association of Anatomists 209(3): 310–322.

Morante, Javier, Vallejo, Diana M, Desplan, Claude and Dominguez, Maria (2013) ‘Conserved miR-8/miR-200 defines a glial niche that controls neuroepithelial expansion and neuroblast transition’, Developmental cell 27(2): 174–187.

Morgan, S. C., Taylor, D. L. and Pocock, J. M. (2004) ‘Microglia release activators of neuronal proliferation mediated by activation of mitogen-activated protein kinase, phosphatidylinositol-3-kinase/Akt and delta-Notch signalling cascades’, J Neurochem 90(1): 89–101.

O’Neill, Elizabeth M, Rebay, Ilaria, Tjian, Robert and Rubin, Gerald M (1994) ’The activities of two Ets-related transcription factors required for Drosophila eye development are modulated by the Ras/MAPK pathway’, Cell 78(1): 137–147.

Ozel, M. N., Simon, F., Jafari, S., Holguera, I., Chen, Y. C., Benhra, N., El-Danaf, R. N., Kapuralin, K., Malin, J. A., Konstantinides, N. et al. (2021) ‘Neuronal diversity and convergence in a visual system developmental atlas’, Nature 589(7840): 88–95.

Phillips, R. G., Roberts, I. J., Ingham, P. W. and Whittle, J. R. (1990) ‘The Drosophila segment polarity gene patched is involved in a position-signalling mechanism in imaginal discs’, Development 110(1): 105–14.

Prasad, A. R., Lago-Baldaia, I., Bostock, M. P., Housseini, Z. and Fernandes, V. M. (2022) ‘Differentiation signals from glia are fine-tuned to set neuronal numbers during development’, Elife 11.

Queenan, Anne Marie, Ghabrial, Amin and Schüpbach, Trudi (1997) ‘Ectopic activation of torpedo/Egfr, a Drosophila receptor tyrosine kinase, dorsalizes both the eggshell and the embryo’, Development 124(19): 3871–3880.

Rebay, Ilaria and Rubin, Gerald M (1995) ‘Yan functions as a general inhibitor of differentiation and is negatively regulated by activation of the Ras1/MAPK pathway’, Cell 81(6): 857–866.

Robinow, S. and White, K. (1991) ‘Characterization and spatial distribution of the ELAV protein during Drosophila melanogaster development’, J Neurobiol 22(5): 443–61.

Sagner, A. and Briscoe, J. (2019) ‘Establishing neuronal diversity in the spinal cord: a time and a place’, Development 146(22).

Sarov, Mihail, Barz, Christiane, Jambor, Helena, Hein, Marco Y, Schmied, Christopher, Suchold, Dana, Stender, Bettina, Janosch, Stephan, Kj, Vinay Vikas and Krishnan, RT (2016) ‘A genome-wide resource for the analysis of protein localisation in Drosophila’, Elife 5: e12068.

Sato, M., Yasugi, T., Minami, Y., Miura, T. and Nagayama, M. (2016) ‘Notch-mediated lateral inhibition regulates proneural wave propagation when combined with EGF-mediated reaction diffusion’, Proc Natl Acad Sci U S A 113(35): E5153–62.

Sawyer, Jessica K, Kabiri, Zahra, Montague, Ruth A, Allen, Scott R, Stewart, Rebeccah, Paramore, Sarah V, Cohen, Erez, Zaribafzadeh, Hamed, Counter, Christopher M and Fox, Donald T (2020) ‘Exploiting codon usage identifies intensity-specific modifiers of Ras/MAPK signaling in vivo’, PLoS Genetics 16(12): e1009228.

Schindelin, J., Arganda-Carreras, I., Frise, E., Kaynig, V., Longair, M., Pietzsch, T., Preibisch, S., Rueden, C., Saalfeld, S., Schmid, B. et al. (2012) ‘Fiji: an open-source platform for biological-image analysis’, Nat Methods 9(7): 676–82.

Selleck, Scott B, Gonzalez, Cayetano, Glover, David M and White, Kalpana (1992) ‘Regulation of the G1-S transition in postembryonic neuronal precursors by axon ingrowth’, Nature 355(6357): 253–255.

Shwartz, A., Yogev, S., Schejter, E. D. and Shilo, B. Z. (2013) ‘Sequential activation of ETS proteins provides a sustained transcriptional response to EGFR signaling’, Development 140(13): 2746–54.

Suzuki, T., Takayama, R. and Sato, M. (2016a) ‘eyeless/Pax6 controls the production of glial cells in the visual center of Drosophila melanogaster’, Dev Biol 409(2): 343–53.

Suzuki, T., Trush, O., Yasugi, T., Takayama, R. and Sato, M. (2016b) ‘Wnt Signaling Specifies Anteroposterior Progenitor Zone Identity in the Drosophila Visual Center’, J Neurosci 36(24): 6503–13.

Thibert, Chantal, Teillet, Marie-Aimée, Lapointe, Françoise, Mazelin, Laetitia, Le Douarin, Nicole M and Mehlen, Patrick (2003) ‘Inhibition of neuroepithelial patched-induced apoptosis by sonic hedgehog’, Science 301(5634): 843–846.

Weng, M., Haenfler, J. M. and Lee, C. Y. (2012) ‘Changes in Notch signaling coordinates maintenance and differentiation of the Drosophila larval optic lobe neuroepithelia’, Dev Neurobiol 72(11): 1376–90.

Wilcockson, S. G., Guglielmi, L., Araguas Rodriguez, P., Amoyel, M. and Hill, C. S. (2023) ‘An improved Erk biosensor detects oscillatory Erk dynamics driven by mitotic erasure during early development’, Dev Cell 58(23): 2802–2818 e5.

Wu, C., Boisclair Lachance, J. F., Ludwig, M. Z. and Rebay, I. (2020) ‘A context-dependent bifurcation in the Pointed transcriptional effector network contributes specificity and robustness to retinal cell fate acquisition’, PLoS Genet 16(11): e1009216.

Yasugi, T., Umetsu, D., Murakami, S., Sato, M. and Tabata, T. (2008) ‘Drosophila optic lobe neuroblasts triggered by a wave of proneural gene expression that is negatively regulated by JAK/STAT’, Development 135(8): 1471–80.

Yasugi, Tetsuo, Sugie, Atsushi, Umetsu, Daiki and Tabata, Tetsuya (2010) ‘Coordinated sequential action of EGFR and Notch signaling pathways regulates proneural wave progression in the Drosophila optic lobe’, Development 137(19): 3193–3203.

Yoshida, S., Soustelle, L., Giangrande, A., Umetsu, D., Murakami, S., Yasugi, T., Awasaki, T., Ito, K., Sato, M. and Tabata, T. (2005) ‘DPP signaling controls development of the lamina glia required for retinal axon targeting in the visual system of Drosophila’, Development 132(20): 4587–98.

Yuen, Alice C, Prasad, Anadika R, Fernandes, Vilaiwan M and Amoyel, Marc (2022) ‘A kinase translocation reporter reveals real-time dynamics of ERK activity in Drosophila’, Biology Open 11(5): bio059364.

