## Supplementary Information for "Glia generate distinct visual processing centres by locally inhibiting ERK activity in an optic lobe neuroepithelium"

**for**

by

Bradleigh M. J. Cocker[1,§], Matthew P. Bostock[1,2,§], Haoruo Wei[1], Vilaiwan M. Fernandes[1]

Affiliations:

1. Cell and Developmental Biology, Faculty of Life Sciences, University College London, London, UK
2. Current Address: Centre for Developmental Neurobiology, Institute of Psychiatry, Psychology & Neuroscience, King's College London, London, UK

[§]These authors contributed equally to this work

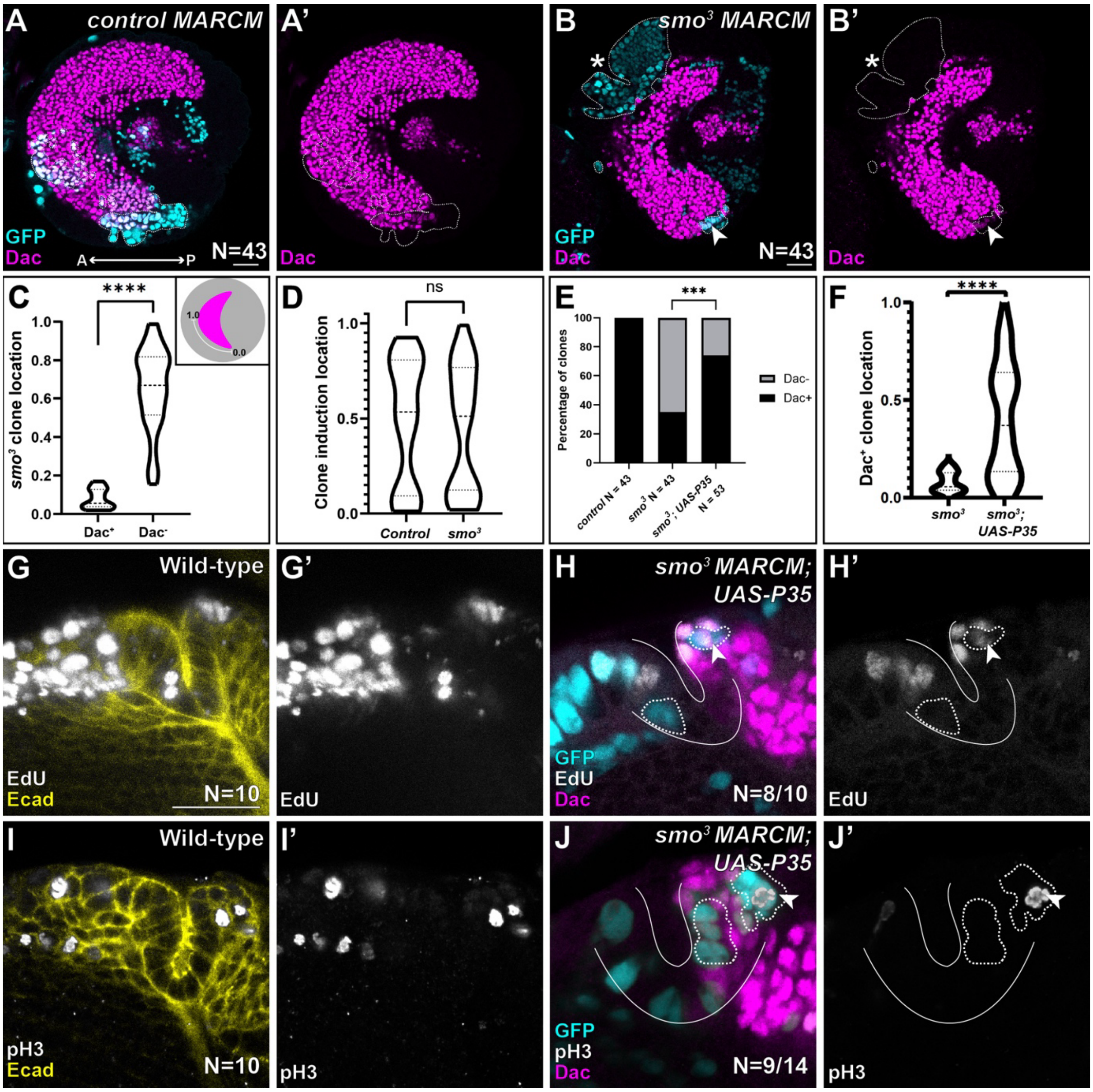


**Fig.S1: Hh signalling is dispensable for Dac expression at the lamina tips and cell cycle progression throughout the lamina.**

**(A-B)** Representative lateral views of optic lobes containing control (A; N=43) and *smo*$^{3}$ (B; N=43) MARCM clones marked with GFP (cyan - outlined) and Dac (magenta).

**(C,D)** Distributions of the positions from the centre (1.0) to the tips (0.0) of the lamina crescent of *smo*$^{3}$ MARCM clones containing Dac+ cells and not containing any Dac expressing cells (Dac-) (unpaired, two-tailed t test, P<0.0001).

**(D)** Distributions of the positions of control and *smo*$^{3}$ MARCM clones (unpaired, two-tailed t test, P=0.6742).

**(E)** Quantification of the percentage of control clones (N=43), *smo*$^{3}$ clones (N=43) and *smo*$^{3}$ clones expressing P35 (N=53) containing cells expressing Dac (Fisher's Exact test P<0.0002).

**(F)** Distributions of the positions of *smo*$^3$ MARCM and *smo*$^3$ MARCM clones expressing P35 (N=53 clones) containing cells expressing Dac (unpaired, Mann Whitney test, P<0.0001).

**(G-J)** Representative cross-sections of wild-type optic lobes (G and I; N = 10) and optic lobes containing *smo*$^3$ MARCM clones expressing P35 (H and J) marked by GFP (cyan - outlined) stained for Dac (magenta), Ecad (G and I; yellow) and the S-phase marker EdU (G and H; grey) or the mitotic marker Phospho-Histone H3 (I andJ; grey). White lines indicate the lamina furrow, arrowheads mark clones containing EdU-positive(H; N=8/10 clones) or pH3-positive (J; N=9/14 clones) cells.

All optic lobes are from pupae dissected at 0h APF. Scale bar = 20 μm. Anterior (A), posterior (P), medial (M), lateral (L).

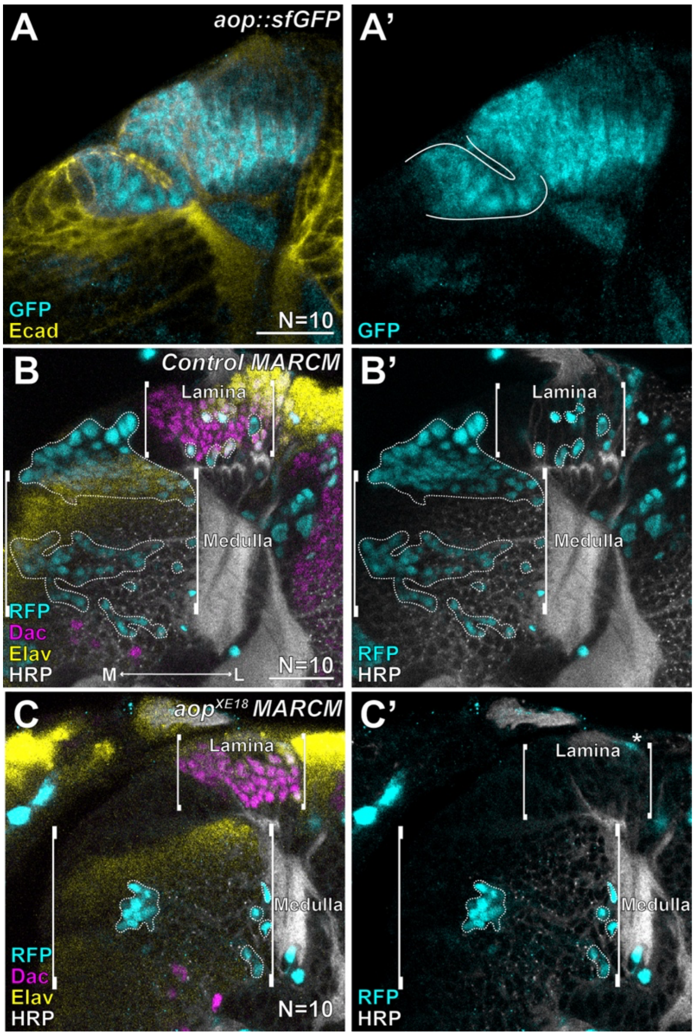


**Fig. S2: Aop, which represses ERK-dependent transcription is required cell autonomously for cells to take on lamina fate.**

**(A)** Representative optic lobe expressing Aop::sfGFP (cyan) and Ecad (yellow). White lines indicate the neuroepithelium.

**(B,C)** Representative cross-sections of optic lobes containing control (A; N=10 lobes) and *aop*$^{XE18}$ (N=10 lobes) MARCM clones marked by RFP (cyan - outlined) stained for Dac (magenta), the pan-neuronal marker Elav (yellow) and HRP (grey). Brackets indicate the approximate regions of the lamina and medulla. Asterisk marks a glial cell.

Scale bar = 20 μm. All optic lobes are from late L3 larvae. medial (M), lateral (L).

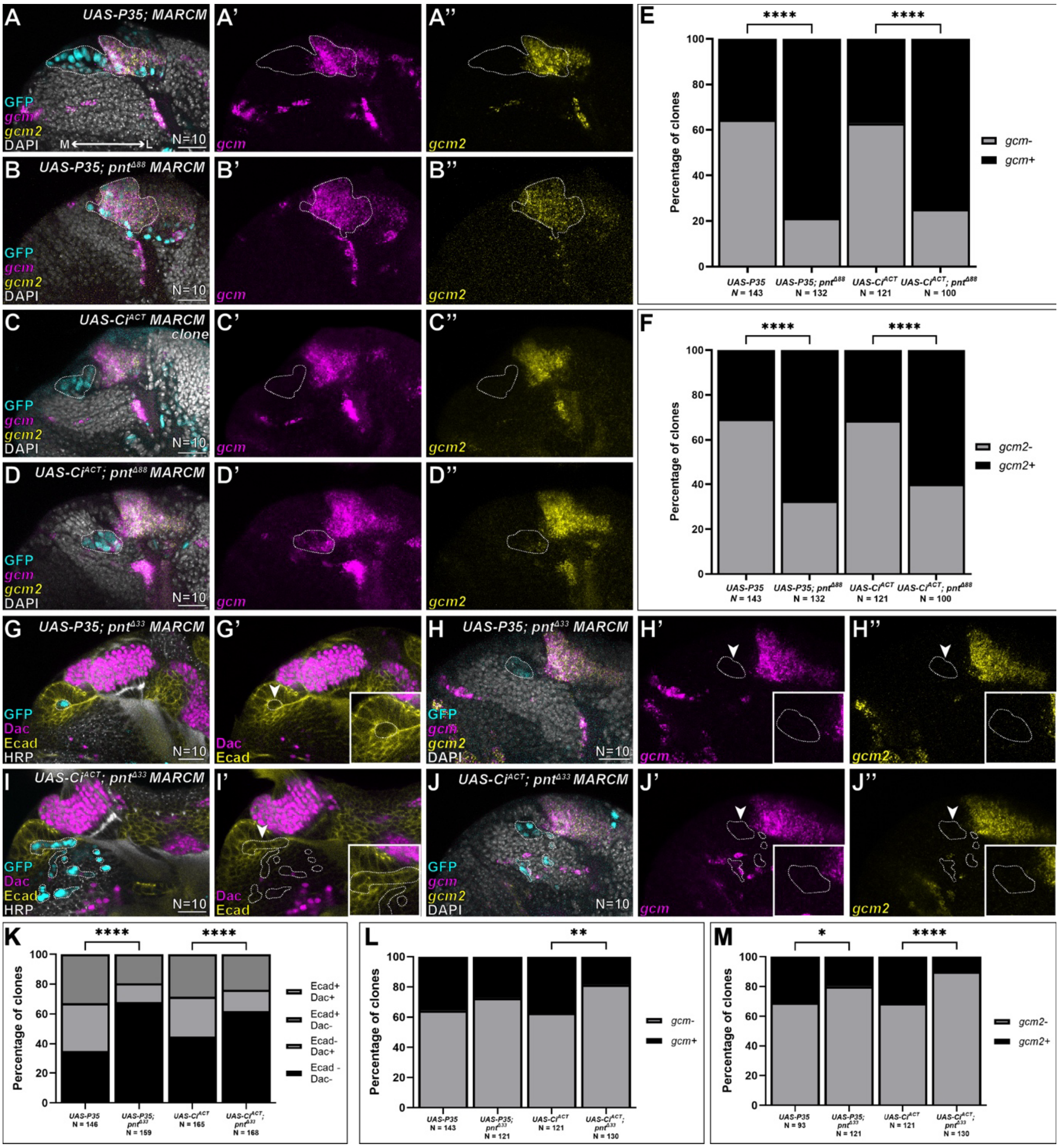

**Fig. S3: *pnt* isoforms cooperate to antagonise lamina fate in the neuroepithelium.**

Representative cross-sections of optic lobes containing MARCM clones marked by GFP (cyan - outlined) stained for *gcm* (magenta), *gcm2* (yellow) and with nuclei stained by DAPI (grey) for the following genotypes:

**(A)** control clones expressing P35

**(B)** $pnt^{\Delta 88}$ mutant clones expressing P35

**(C)** control clones expressing $Ci^{ACT}$

**(D)** $pnt^{\Delta 88}$ mutant clones expressing $Ci^{ACT}$

*gcm* and *gcm2* are expressed in LPCs as well as in glia in the medulla cortex.

**(E,F)** Quantifications of the percentage of clones from (A-D) containing cells medial to the lamina furrow expressing *gcm* and *gcm2* (Fisher's Exact tests, **** = P<0.0001).
Representative cross-sections of optic lobes containing MARCM marked by GFP (cyan - outlined) stained for either Dac (magenta), Ecad (yellow) and HRP (grey) in (G,I) or *gcm* (magenta), *gcm2* (yellow) and DAPI (grey) (H,J). Insets show magnified images of clones indicated by white arrowheads for the following genotypes:
**(G, H)** $pnt^{\Delta 33}$ MARCM clones expressing P35
**(I,J)** $pnt^{\Delta 33}$ MARCM clones expressing $Ci^{ACT}$
**(K)** Quantification of the percentage of clones from (G,I) containing cells expressing Dac and Ecad (Fisher's Exact test, **** = P<0.0001).
**(L,M)** Quantifications of the percentage of clones from (H,J) containing cells expressing *gcm* (L) or *gcm2* (M) (Fisher's exact tests, * = P = 0.0485, ** = P = 0.0011, **** = P<0.0001).
Scale bar = 20 μm. Medial (M), lateral (L).

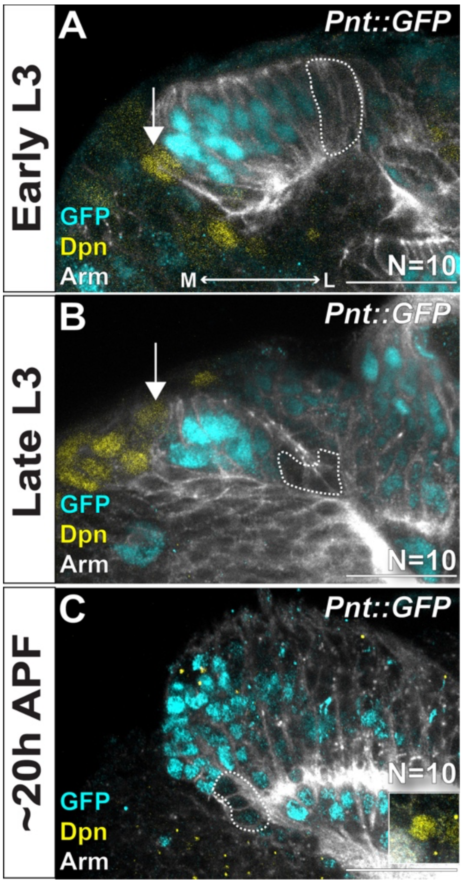


**Fig. S4: ERK signalling is absent from the lateral neuroepithelium up to medulla neuroblast termination.**

Time series of representative cross-sections from Pnt::GFP (cyan) optic lobes stained for Dpn (yellow) and Arm (grey) from:

**(A)** early L3 (N=10 lobes)

**(B)** late L3 (N=10 lobes)

**(C)** ~20 hours APF (N=10 lobes).

Dashed lines outline the absence of Pnt::GFP expression at the lateral neuroepithelium, arrows indicate the youngest medulla neuroblast. Inset in (C) shows expression of Dpn within the developing lobula complex of the same optic lobe.

Scale bar = 20 μm. Medial (M), lateral (L).

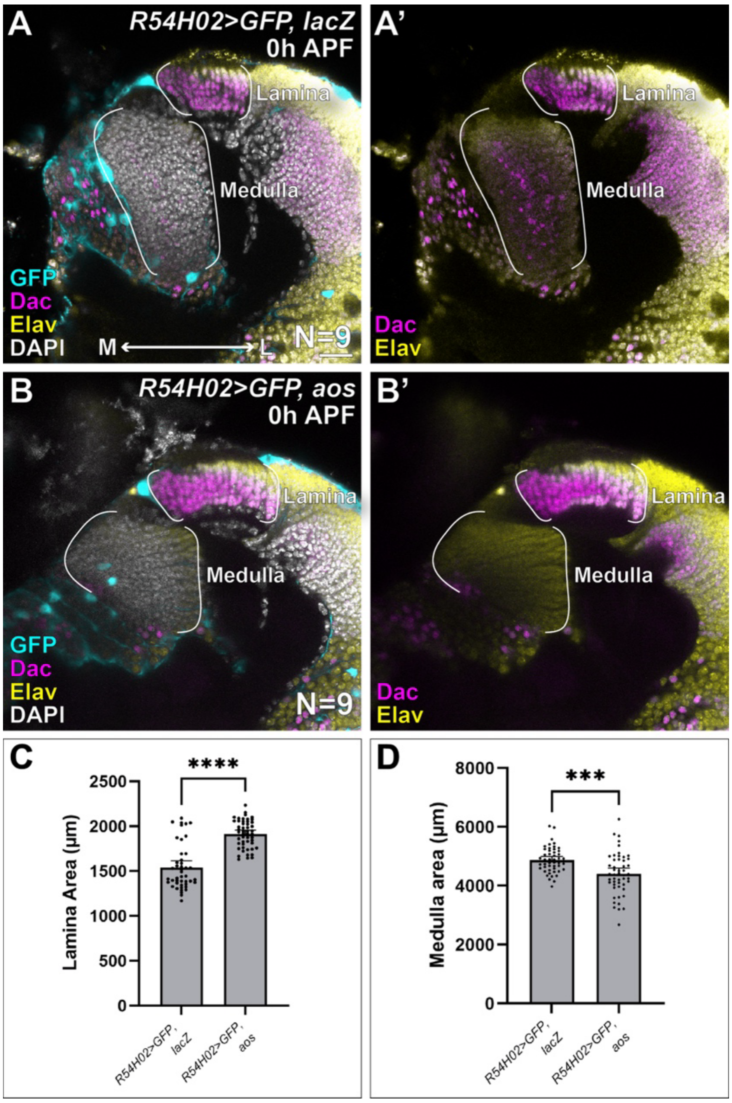


**Fig. S5: Overexpression of *aos* in cortex glia increases the size of the lamina at expense of the medulla.**

**(A,B)** Representative optic lobes from pupae at 0h APF with R54H02-Gal4 driving expression of GFP and *lacZ* (A; N=9) or GFP and *aos* (B; N=9) in cortex glia. Optic lobes were stained for GFP (cyan), Dac (magenta), Elav (yellow) and DAPI (grey). White lines indicate the approximate area of the medulla and lamina.

**(C,D)** Quantifications of lamina and medulla area (unpaired t tests, **** = P<0.0001, *** = P=0.0001). Scale bar = 20 µm. Anterior (A), posterior (P), medial (M), lateral (L).

**Table S1: Full experimental genotypes and experimental conditions associated with each figure panel.** (Note that only female genotypes are listed though both sexes were included in our analyses)

| **Figure** | **Panel** | **Genotype** | **Conditions** |
|---|---|---|---|
| 1 | A,B | *;Ecad::GFP;* | Raised at 25°C |
| 1 | C | *;Ptc::GFP;* | Raised at 25°C |
| 2 | A,D,F | Wild-type (Canton S) | Raised at 25°C |
| 2 | B | *y,w,hsflp122/+, tub-Gal4, UAS-nls.GFP; FRT42D, tub-Gal80/FRT42D,* $ptc^{S2}$*;* | Raised at 25°C before and after heat shocking. See Mosaic analysis for more detail. |
| 2 | E,G | *y,w,hsflp122/+, tub-Gal4, UAS-nls.GFP; FRT40A, tub-Gal80/FRT40A,* $smo^{3}$*;* | Raised at 25°C before and after heat shocking. See Mosaic analysis for more detail. |
| 2 | H | *y,w,hsflp122/+, tub-Gal4, UAS-nls.GFP; FRT40A, tub-Gal80/FRT40A,* $smo^{3}$*; UAS-P35/+* | Raised at 25°C before and after heat shocking. See Mosaic analysis for more detail. |
| S1 | A | *y,w,hsflp122/+, tub-Gal4, UAS-nls.GFP; FRT40A, tub-Gal80/FRT40A;* | Raised at 25°C before and after heat shocking. See Mosaic analysis for more detail. |
| S1 | B | *y,w,hsflp122/+, tub-Gal4, UAS-nls.GFP; FRT40A, tub-Gal80/FRT40A,* $smo^{3}$*;* | Raised at 25°C before and after heat shocking. See Mosaic analysis for more detail. |
| S1 | G,I | Wild-type (Canton S) | Raised at 25°C |
| S1 | H,J | *y,w,hsflp122/+, tub-Gal4, UAS-nls.GFP; FRT40A, tub-Gal80/FRT40A,* $smo^{3}$*; UAS-P35/+* | Raised at 25°C before and after heat shocking. See Mosaic analysis for more detail. |
| 3 | A,B,C | *y,w,*$hsflp^{122}$/+; *UAS-CD8::GFP/+*; *pxb-T2AGal4/+* | Raised at 25°C for 7 days AEL, at which point they were shifted to 29°C for 48 hours |

| | | | |
|---|---|---|---|
| 3 | D,E,F | *y,w,hsflp$^{122}$/+; UAS-CD8::GFP/+; pxb-T2AGal4/UAS-EGFR$^{ACT}$* | Raised at 25°C for 7 days AEL, at which point they were shifted to 29°C for 48 hours |
| 3 | G,H,I | *y,w,hsflp$^{122}$/ +; UAS-CD8::GFP/+; pxb-T2AGal4/UAS-aop$^{RNAi}$* | Raised at 25°C for 7 days AEL, at which point they were shifted to 29°C for 48 hours |
| S2 | A | *;;aop::sfGFP* | Raised at 25°C |
| S2 | B | *y,w,hsflp$^{122}$/w, Tub>Gal4, UAS-mCherry.nls; Tub>Gal80, FRT40A/FRT40A;* | Raised at 25°C before and after heat shocking. See Mosaic analysis for more detail. |
| S2 | C | *y,w,hsflp$^{122}$/w, Tub>Gal4, UAS-mCherry.nls; Tub>Gal80, FRT40A/aop$^{EX18}$, FRT40A* | Raised at 25°C before and after heat shocking. See Mosaic analysis for more detail. |
| 4 | A,C | *y,w,hsflp$^{122}$, Tub>Gal4, UAS-nls.GFP/ y,w,hsflp$^{122}$;; Tub>Gal80, FRT80B/Dronc$^{I24}$, FRT80B* | Raised at 25°C before and after heat shocking. See Mosaic analysis for more detail. |
| 4 | B,D | *y,w,hsflp$^{122}$, Tub>Gal4, UAS-nls.GFP/ y,w,hsflp$^{122}$;UAS-aop$^{ACT}$/+; Tub>Gal80, FRT80B/Dronc$^{I24}$, FRT80B* | Raised at 25°C before and after heat shocking. See Mosaic analysis for more detail. |
| 4 | E | *y,w,hsflp$^{122}$, Tub>Gal4, UAS-nls.GFP; UAS-P35/+; Tub>Gal80, FRT82B/ FRT82B* | Raised at 25°C before and after heat shocking. See Mosaic analysis for more detail. |
| 4 | F | *y,w,hsflp$^{122}$, Tub>Gal4, UAS-nls.GFP; UAS-P35/+; Tub>Gal80, FRT82B/ pnt$^{\Delta 88}$, FRT82B* | Raised at 25°C before and after heat shocking. See Mosaic analysis for more detail. |
| 4 | G | *y,w,hsflp$^{122}$, Tub>Gal4, UAS-nls.GFP; UAS-ci$^{ACT}$/+; Tub>Gal80, FRT82B/ FRT82B* | Raised at 25°C before and after heat shocking. See Mosaic analysis for more detail. |

| | | | |
|---|---|---|---|
| 4 | H | *y,w,hsflp$^{122}$, Tub>Gal4, UAS-nls.GFP; UAS-ci$^{ACT}$/+; Tub>Gal80, FRT82B/ pnt$^{\Delta 88}$, FRT82B* | Raised at 25°C before and after heat shocking. See Mosaic analysis for more detail. |
| S3 | A | *y,w,hsflp$^{122}$, Tub>Gal4, UAS-nls.GFP; UAS-P35/+; Tub>Gal80, FRT82B/ FRT82B* | Raised at 25°C before and after heat shocking. See Mosaic analysis for more detail. |
| S3 | B | *y,w,hsflp$^{122}$, Tub>Gal4, UAS-nls.GFP; UAS-P35/+; Tub>Gal80, FRT82B/ pnt$^{\Delta 88}$, FRT82B* | Raised at 25°C before and after heat shocking. See Mosaic analysis for more detail. |
| S3 | C | *y,w,hsflp$^{122}$, Tub>Gal4, UAS-nls.GFP; UAS-ci$^{ACT}$/+; Tub>Gal80, FRT82B/ pnt$^{\Delta 88}$, FRT82B* | Raised at 25°C before and after heat shocking. See Mosaic analysis for more detail. |
| S3 | D | *y,w,hsflp$^{122}$, Tub>Gal4, UAS-nls.GFP; UAS-ci$^{ACT}$/+; Tub>Gal80, FRT82B/ pnt$^{\Delta 88}$, FRT82B* | Raised at 25°C before and after heat shocking. See Mosaic analysis for more detail. |
| S3 | G,H | *y,w,hsflp$^{122}$, Tub>Gal4, UAS-nls.GFP; UAS-P35/+; Tub>Gal80, FRT82B/ pnt$^{\Delta 33}$, FRT82B* | Raised at 25°C before and after heat shocking. See Mosaic analysis for more detail. |
| S3 | I,J | *y,w,hsflp$^{122}$, Tub>Gal4, UAS-nls.GFP; UAS-ci$^{ACT}$/+; Tub>Gal80, FRT82B/ pnt$^{\Delta 33}$, FRT82B* | Raised at 25°C before and after heat shocking. See Mosaic analysis for more detail. |
| 5 | A,C,E,G | *; Pnt::GFP;* | Raised at 25°C |
| 5 | I | *w$^{1118}$/y$^{1}$,w$^{*}$; UAS-modERK-KTR-T2A-His2AV-mCh/+; C855a-Gal4/+* | Raised at 25°C |
| S4 | A,B,C | *; Pnt::GFP;* | Raised at 25°C |
| 6 | A | *w$^{1118}$/y$^{1}$,w$^{*}$; UAS-GFP.nls/UAS-nls.LacZ; R54H02-Gal4/UAS-nls.LacZ* | Raised at 25°C |
| 6 | B,D,F | *w/+;; aos-LacZ$^{w11}$/+* | Raised at 25°C |
| 6 | C | *;; aos:sfGFP:* | Raised at 25°C |
| 6 | E,G | *w/+;; aos-LacZ$^{w11}$/aos$^{\Delta 7}$* | Raised at 25°C |

| | | | |
|---|---|---|---|
| 6 | H | *$y^1$,$w^*$/$w^*$; wrapper-Gal4/UAS-nls.LacZ; aos-LacZ$^{w11}$/aos$^{\Delta 7}$* | Raised at 25°C |
| 6 | I | *$y^1$,$w^*$/$w^*$; wrapper-Gal4/UAS-aos; aos-LacZ$^{w11}$/aos$^{\Delta 7}$* | Raised at 25°C |
| S4 | A | *y,w,hsflp$^{122}$; UAS-nls.GFP/UAS-nls.LacZ; R54H02-Gal4/UAS-nls.LacZ* | Raised at 25°C |
| S4 | B | *y,w,hsflp$^{122}$; UAS-nls.GFP/UAS-aos; R54H02-Gal4/UAS-aos* | Raised at 25°C |

**Table S2: probe sequences for *gcm* and *gcm2***

| ***gcm*** | ***gcm2*** |
|---|---|
| GTCCCTGCCTCTATATCTTTCGACTGCGCAAGGCTGACTAACTGG | CTCACTCCCAATCTCTATAACCGATGACTCGGATCCACTTAGTGG |
| TCTCTGGAGCGACGCGTCCGTTCGCTTCCACTCAACTTTAACCCG | ACACATTAGCTGCCGTCTCGCAGGGAACTACCCTACAAATCCAAT |
| GTCCCTGCCTCTATATCTTTAGTGGAGCATATCGTAGGAGTACAT | CTCACTCCCAATCTCTATAACTAGTCTACCGCCACGACAACTTTT |
| CCGTGGAGCTATGCGCTGAGAGAGGTTCCACTCAACTTTAACCCG | CGCAGTGACCGCGACACGGTTTAATAACTACCCTACAAATCCAAT |
| GTCCCTGCCTCTATATCTTTAAGGCTGCTTCGATTAAGTTTTTCT | CTCACTCCCAATCTCTATAATTTTCGCTGAACCTCGAATACAACC |
| TGCTTGACACGTGTCAATTCTCAAATTCCACTCAACTTTAACCCG | TGGGAAACAACCGTGAGAATGACGAAACTACCCTACAAATCCAAT |
| GTCCCTGCCTCTATATCTTTGGCCGCGGCAAGCCTGGATTTCCAA | CTCACTCCCAATCTCTATAATTCAATGGGTGACCGATGGGGCTAG |
| AGAAGTGGGTCACTGGATAGCCGCATTCCACTCAACTTTAACCCG | TCCGGTGTCCGGATTGGTTAAGTACAACTACCCTACAAATCCAAT |
| GTCCCTGCCTCTATATCTTTTGGTGCTATGTGTGGGCGTCGATGT | CTCACTCCCAATCTCTATAATAGTTCCTGTGTCGAAATAGCTGAC |
| CATATGAACCCTGAGATTGATTAAATTCCACTCAACTTTAACCCG | CGGTTGCAGTATTTCCTGGTGCCGTAACTACCCTACAAATCCAAT |
| GTCCCTGCCTCTATATCTTTTCTGCACGGAGTAGATGAGCCGGCA | CTCACTCCCAATCTCTATAATCGCGGATAGCCGTCGCTAGTTGAA |
| CGCTGGCATGTTTCCTGGCCTCATCTTCCACTCAACTTTAACCCG | CACTTGTATCCACTGGGCACTTGTAAACTACCCTACAAATCCAAT |
| GTCCCTGCCTCTATATCTTTTGGTTGTGTAGAAACTTGCATCCTG | CTCACTCCCAATCTCTATAATGGAATCATAGGCTACTGCCTGGCA |
| ACTCAAACACACTGAAGATCTCCGATTCCACTCAACTTTAACCCG | CCGGTTCCGGAGTGGCCAGTTGCGGAACTACCCTACAAATCCAAT |
| GTCCCTGCCTCTATATCTTTTGTCGCTCAGCGCCGATTCCCTGGC | CTCACTCCCAATCTCTATAATGCTGTCATCCAGGTAGCTGCAAGT |
| GTTTTGTGGGTCTCAGGCTACTCAGTTCCACTCAACTTTAACCCG | GTGACAACTGATGGCTCCTCGCCGTAACTACCCTACAAATCCAAT |
| GTCCCTGCCTCTATATCTTTTGTTGACTGGGTGATATGTCGCGTC | CTCACTCCCAATCTCTATAACCAGCGGCACACGTCCAAAGGCCCG |
| TCCCCTAGCAATAGATGGGATCCGTTTCCACTCAACTTTAACCCG | CGACTGATCCGTTGGCTGACTTTCCAACTACCCTACAAATCCAAT |
| GTCCCTGCCTCTATATCTTTTAGTTGCGATGAGAGATCTTATCCC | CTCACTCCCAATCTCTATAAGTTGCTGCGTCTTGAAGCTGTAGCC |
| AACAACGCTTGAGTGGCTCAGTAATTTCCACTCAACTTTAACCCG | GAGAATGATCGGGTTGCTGTGTGAGAACTACCCTACAAATCCAAT |
| GTCCCTGCCTCTATATCTTTCACTGGCTCCATTGGGCAACTTGCA | CTCACTCCCAATCTCTATAAAGTGATGTCCCGCTGTCTGATTTCC |
| TGTCGCAAATGGCGGGTCTCAGATGTTCCACTCAACTTTAACCCG | CATTATAGCCACTACTGCTCTCGTAAACTACCCTACAAATCCAAT |
| GTCCCTGCCTCTATATCTTTGCATTCCGCACTGAAGGCTGGAGGA | CTCACTCCCAATCTCTATAATGCAGGCGTAGGGCAAACTGGGAGA |
| ACGTATCGTCGCATATTTCATACGATTCCACTCAACTTTAACCCG | GCTGGTATGCGGCCAGTTCCGAAATAACTACCCTACAAATCCAAT |
| GTCCCTGCCTCTATATCTTTTTACACGATTTGGTTGACTAACGA | CTCACTCCCAATCTCTATAACGTAGCTGGTGTAGTGATCGTAGTA |

| | |
|---|---|
| ATTTTCTAGGGCGACAAGCACCCATTTCCACTCAACTTTAACCCG | ATCCACCTGCTACCGCCATTTGTTGAACTACCCTACAAATCCAAT |
| GTCCCTGCCTCTATATCTTTCGTAGTGAATCTCTCTATTCCACTG | CTCACTCCCAATCTCTATAAGCAGGTTGATCTTGGATCCGTTGGG |
| CATTGGCATAGCATGGATCCTCCGTTTCCACTCAACTTTAACCCG | GCCGGGCTTTGTCGCAAATGGCTGGAACTACCCTACAAATCCAAT |
| GTCCCTGCCTCTATATCTTTAAACCATTTTTGATGTGTCTATGTA | CTCACTCCCAATCTCTATAAGCCGCCAGAAGTGGGTCACCGGATA |
| CTGGCATTGTTATAGGCATGCCGTTTTCCACTCAACTTTAACCCG | CCTGAAAAAAGATGGCATTGCCGGAAACTACCCTACAAATCCAAT |
| GTCCCTGCCTCTATATCTTTGCTGGTATTGTGAGTGCTGCAGTTG | CTCACTCCCAATCTCTATAAGACGCAGGTGATCATGCACTCCCTT |
| GTACATGTTGCTGCGGTGAGAGTTGTTCCACTCAACTTTAACCCG | TGGAAACACTGGAGTTCTTCGGATCAACTACCCTACAAATCCAAT |
| GTCCCTGCCTCTATATCTTTTTCCTCCTGCCAAAAGTCGCCTGGC | CTCACTCCCAATCTCTATAACCGGATGCACCGAATGCGTACTCTG |
| ACATTACGGCCAAACTTCGCACACGTTCCACTCAACTTTAACCCG | CCGCCTGCTGTTGACCAGGACTTGGAACTACCCTACAAATCCAAT |
| GTCCCTGCCTCTATATCTTTGGTGATCGTGGGTGCCCTTGGCCTG | CTCACTCCCAATCTCTATAACCCACTCCCGTTTTCCCTTGCCCGT |
| CCGTGGATCCCTTGGCTTCGGGTCTTTCCACTCAACTTTAACCCG | CATGTGGCACAATGGCATCATTGATAACTACCCTACAAATCCAAT |
| GTCCCTGCCTCTATATCTTTAACTAGAAACGTTCAATATATAATA | CTCACTCCCAATCTCTATAAAGGCATTATACTGATATATATCCAA |
| TAGCTGGGAAACAAGTACAAAAGTATTCCACTCAACTTTAACCCG | AGTGGCTATAACCGGCGCACTTGCCAACTACCCTACAAATCCAAT |
| GTCCCTGCCTCTATATCTTTTACTACTGTGTACTTCTTCTCAACT | CTCACTCCCAATCTCTATAACTTGAGCATTGTAATCCAATTAGGC |
| CTATAGTTTTCCTTCTCAATAGTCTTTCCACTCAACTTTAACCCG | ATTCGAAAGCCCTGCGAACAGGCAAAACTACCCTACAAATCCAAT |
